# Live-cell imaging shows uneven segregation of extrachromosomal DNA elements and transcriptionally active extrachromosomal DNA clusters in cancer

**DOI:** 10.1101/2020.10.20.335216

**Authors:** Eunhee Yi, Amit D. Gujar, Molly Guthrie, Hoon Kim, Kevin C. Johnson, Samirkumar B. Amin, Sunit Das, Patricia A. Clow, Albert W. Cheng, Roel GW Verhaak

## Abstract

Oncogenic extrachromosomal DNA elements (ecDNAs) promote intratumoral heterogeneity, creating a barrier for successful cancer treatments. The underlying mechanisms are poorly understood and studies are hampered in part by a lack of adequate tools enabling studies of ecDNA behavior. Here, we show that single-cell ecDNA copy numbers follow a Gaussian distribution across tumor cells in vitro and in patient glioblastoma specimens, suggesting uneven ecDNA segregation during mitosis. We established a CRISPR-based approach which leverages unique ecDNA breakpoint sequences to tag ecDNA with fluorescent markers in living cells. Applying this method during mitosis revealed disjointed ecDNA inheritance patterns, providing an explanation for rapid ecDNA accumulation in cancer. Post-mitosis, ecDNAs tended to cluster and clustered ecDNAs colocalized with RNA polymerase II, promoting transcription of cargo oncogenes. Our observations provide direct evidence for uneven segregation of ecDNA and shed new lights of mechanisms through which ecDNAs contribute to oncogenesis.

## Introduction

Tumor evolution drives intratumor heterogeneity which is a source of therapy failure and resistance^1, 2^. Genomic instability and chromosomal structural variations including oncogene amplification play a critical role in driving tumor evolution^3, 4^. Focal amplifications in cancer may occur on extrachromosomal DNA (ecDNA) elements, are observed in the majority of glioblastomas and at high frequencies in many cancer types ^5–8^, and contribute to an accelerated tumor growth and poor patient survival. EcDNAs are 50kb-5Mb genomic elements containing genes and regulatory sequences^9^. Acentromeric and atelomeric features of ecDNAs suggest uneven ecDNA segregation during mitosis leading to discordant ecDNA inheritance and promoting rapid ecDNA accumulation in a subpopulation of cancer cells. However, direct evidence supporting this hypothesis is lacking^5, 7, 10^. There is also a limited knowledge of ecDNA behavior during DNA replication and our understanding of the ecDNA mobility in proliferating cancer cells. Fluorescence in situ hybridization (FISH) in and 4′,6-diamidino-2-phenylindole (DAPI) staining of fixed cells have been used to demonstrate that oncogenes can reside extrachromosomally in cancer^5, 6, 8 9^. These static readouts of the number of ecDNA copies in single cells are unable to record behavioral patterns. Genome engineering technologies based on the CRISPR-associated RNA-guided inactive endonuclease Cas9 have been leveraged to visualize DNA in living cells. In recent studies, this technique allowed real-time tracing of the dynamic reorganization of genomic DNA during during mitosis^11^, programmable 3D genome interactions^12^ and chromosome translocation induced by genome editing^13^. Although originally designed for labeling of repetitive sequence ^11^, recent advances in CRISPR-based live-cell imaging techniques have additionally enabled visualization of non-repeatitive chromosome loci^14, 15^. These techniques collectively help advance our ability to visualize genome organization during both, physiological and pathological cell states, and in response to CRISPR-guided perturbations. ecDNA sequences are indistinguishable from their parental chromosomal DNA, barring ecDNA-specific breakpoint sequences which provide an opportunity for live-cell ecDNA imaging. Here, we report a CRISPR-based DNA tracking system that leverages DNA breakpoint junctions to label ecDNA elements with multiple fluorescent molecules. We applied this technology to understand ecDNA spatiotemporal dynamics and the mechanisms by which ecDNA contributes to intratumoral heterogeneity.

## Results

### EcDNA shows increased intratumoral copy number variability

While ecDNAhas been nominated as a key factor contributing to intratumoral heterogeneity resulting from suspected unequal segregation ^5, 7, 16^, there is a paucity of direct, experimental evidence supporting this assumption. The proposed model of ecDNA inheritance, in contrast to canonical inheritance of linearly amplified DNA on chromosomes, can account for a high degree of intratumoral multiplicity of ecDNA copy number (Fig. 1A). We hypothesized that the number of ecDNA copies across single cells would be highly variable, whereas the copy number of genes amplified linearly on chromosomes is expected to be identical at the single-cell level. To evaluate the distribution of the number of ecDNA copies per cell, we performed interphase fluorescent in-situ hybridization (FISH) on four glioblastoma (GBM) tumor tissue samples (SMJL006, SMJL012, SMJL017 and SMJL018) and a pair of primary and recurrent GBM neurosphere lines, derived from the same patient (HF3016 and HF3177). We have previously found the GBM oncogene *EGFR* to be focally amplified in all four GBMs and both neurosphere lines^8, 17^. As a control, we included probes mapping to chromosome 7, which was broadly amplified at a low level in all six specimens. We observed a Gaussian distribution pattern of the copy number of *EGFR*-containing ecDNA, demonstrating that the number of ecDNA copies varied widely across cells and implicating uneven segregation of ecDNA (Fig. 1B-C). The copy numbers of chromosome 7 appeared more evenly distributed (Fig. 1B-C, lower panel) but not stable, possibly as a result of cells residing in different stages of the cell cycle and noise levels of the assay. We calculated the median absolute deviation (MAD) of the distribution of copy numbers, which is a metric that represents variability independent of copy number level, and found the MAD of EGFR-ecDNA copies to be significantly higher than the MAD of chromosome 7 copies (average MAD 10.25 vs 1.61; Fligner-Killeen test, p-value < 1.5e-08 in all samples). Representative images of *EGFR*-containing ecDNA copies in cells containing identical numbers of chromosome 7 reflect the impact of ecDNA on intratumoral heterogeneity (Fig. 1B-C, upper panel).

**Fig. 1.**
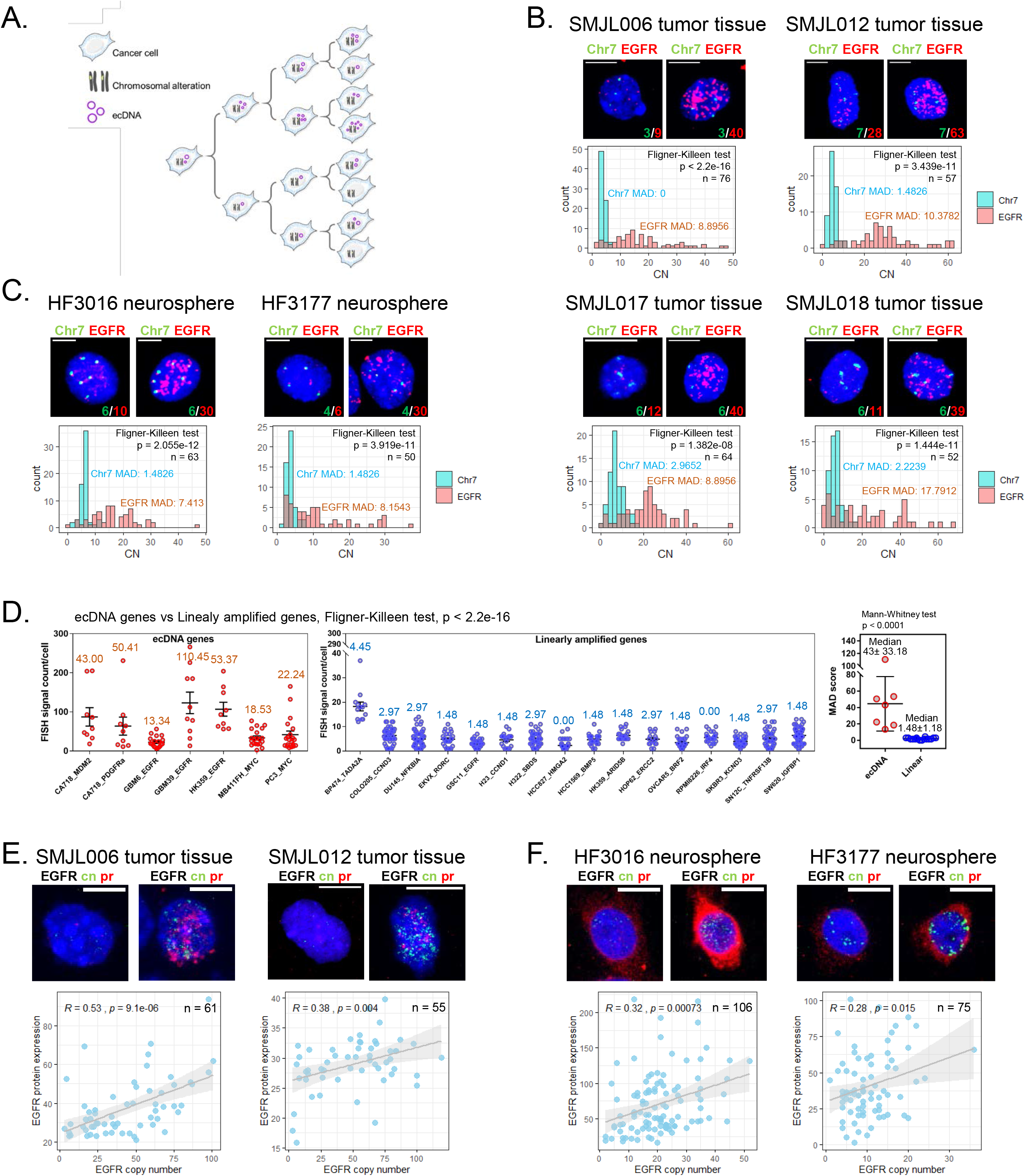
Unevenly segregated ecDNA drives intratumoral heterogeneity. **A.** Cartoon of inheritance pattern of chromosomal alteration and ecDNA. **B-C.** Representative EGFR/Chr7 FISH on four GBM tumor tissues (B, upper panel) and two neurosphere lines (C, upper panel). The MADs are indicated with the corresponding color in each image. Scale bar, 10 μm. Copy number count of each FISH probe per cell and *p* values indicating the homogeneity of variances between EGFR and Chr7 were determined by Fligner-Killeen test (lower panel). **D.** Copy number distribution of ecDNA genes (left panel) and linearly amplified genes (middle panel). The MADs indicated at the top of individual group. *p* value indicating the homogeneity of variances between ecDNA genes and linearly amplified genes was determined by Fligner-Killeen test. The median MAD of ecDNA genes was significantly higher than the median MAD of linearly amplified genes. *p* value indicating significant differences between two group was tested by Mann-Whitney U test. **E-F.** Representative ImmunoFISH experiment on two GBM tumor tissues (E, upper panel) and two neurosphere lines (F, upper panel). Scale bar, 10 μm. Correlation between copy number of EGFR and its protein expression per cell and *p* values were determined by Pearson’s correlation test (lower panel).

To expand our observation, we assessed FISH images across a collection of cell lines from different types and of genes that were recently shown to reside on either linear or circular amplicons by whole-genome sequencing^9^. In total, we compared FISH signals from seven genes on ecDNA and 16 linearly amplified genes. While we observed considerable variability in single-cell copy number of both linear and ecDNA amplicons, ecDNA MADs (median MAD 43 +/− 33.18) were significantly higher than linear amplicon MADs (median MAD 1.48 +/− 1.18; Fig. 1D and Supplementary Fig. 1). The difference in copy number distribution between ecDNA and linear amplicons corroborates previous circumstantial evidence that ecDNA segregates unevenly ^7, 18^.

Intratumoral heterogeneity, which impairs treatment response, is marked by genomic variability as well as intercellular diversity in gene and protein expression^19, 20^. To examine whether the aptitude of ecDNA for enhancing intratumoral diversity ultimately affects the heterogeneity of functional protein expression, we determined the association of *EGFR* copy number with EGFR protein expression at the single-cell level (Fig. 1E-F). In all samples, we observed a positive correlation between *EGFR*-containing ecDNA copy number and EGFR protein expression (Fig. 1E-F, lower panel). As expected, a higher number of ecDNA copies resulted in higher oncogene protein expression in cell line models and patient specimens.

### CRISPR-based labeling enables live-cell ecDNA tracking

To understand how ecDNA heterogeneity is derived, we developed a CRISPR-based DNA labeling method to study ecDNAs in live cells. ecDNAs are formed by DNA breakage followed by end-to-end ligation of DNA segments, resulting in one or more ecDNA breakpoint junctions (Fig. 2A). The sequences covering the breakpoint sites are unique and cannot be detected in the parental linear chromosomes. EcDNA-specific breakpoint sequences provide an opportunity for the design of single guide-RNAs (sgRNA) to label or target ecDNA. We employed Casilio^15, 21^, a hybrid technique that combines dead-Cas9 (dCas) labeling and Pumilio RNA-binding, to recruit multiple fluorescent protein molecules at a prespecified sgRNA target locus (Fig. 2A). SgRNAs designed with programmable Pumilio/FBF (PUF) RNA-binding sites (PUFBSs) achieve target DNA binding through a spacer sequence mapping the ecDNA breakpoint and also enable recruitment of fluorescent molecules conjugated with PUF (Fig. 2A, right panel). We evaluated this approach to label ecDNA-specific breakpoints. To find targetable breakpoint sequences, we analyzed whole-genome sequencing (WGS) data of the HF3016 neurosphere line to reconstruct the structure of the ecDNA element^22^ (Supplementary Fig. 2A-D). This identified four unique ecDNA structures, harboring an *EGFR* fragment (exon1), the full *EGFR* coding sequence, and the non-coding genes *CCAT* and *CCDC26*. We labeled these four lesions as ecEGFRx1, ecEGFR, ecCCAT1 and ecCCDC26 respectively. We designed four primer pairs (Supplementary Fig. 2D, yellow arrows) that allowed extraction of an ecDNA breakpoint fragment from each ecDNA on agarose gels (Supplementary Fig. 2E). We then performed Sanger sequencing to define the precise breakpoint sequence (Supplementary Fig. 2F)^23^.

**Fig. 2.**
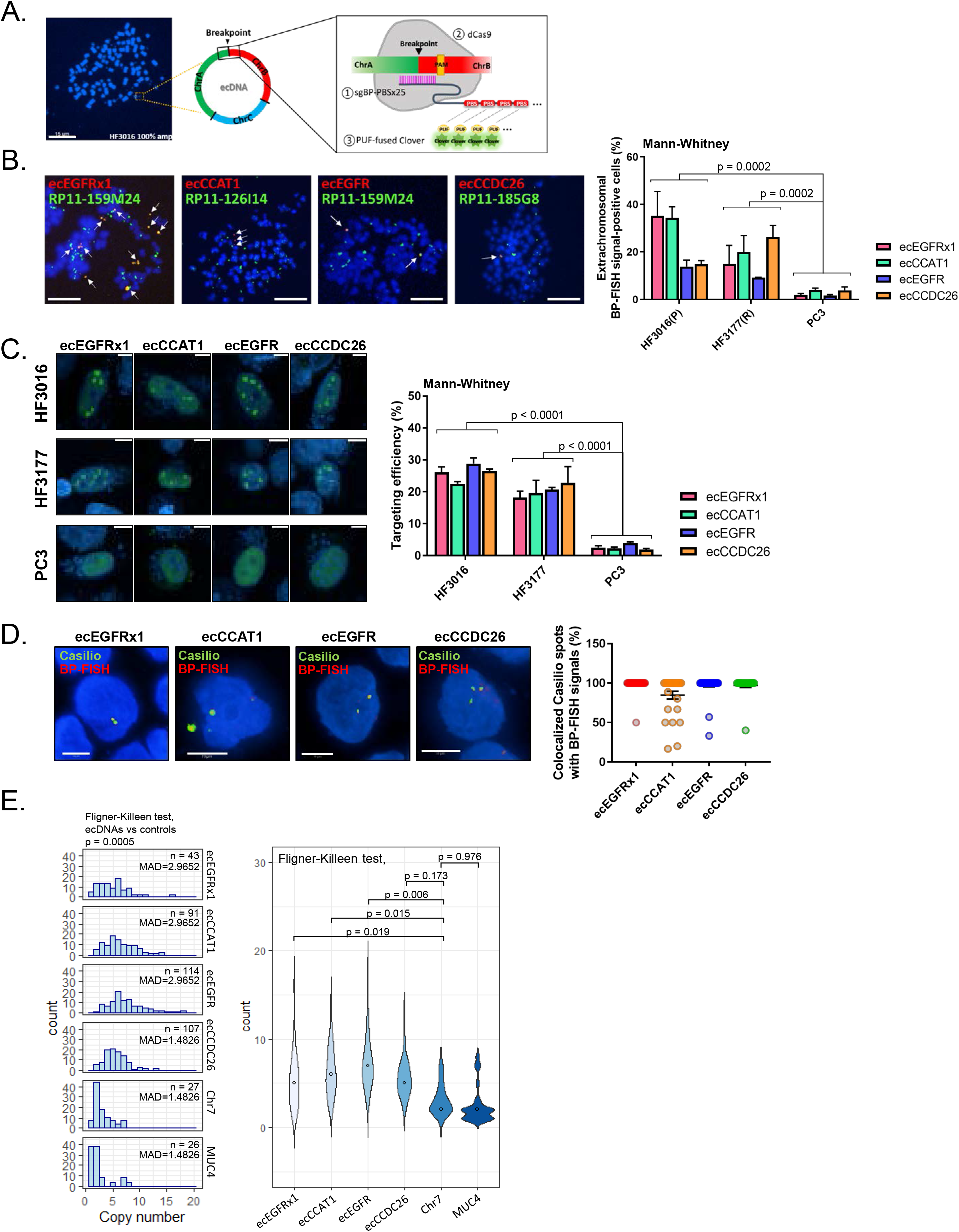
CRISPR-based labeling enables live-cell ecDNA tracking. **A.** Schematic strategy of Casilio-based ecDNA labeling system. **B.** Representative BP-FISH in HF3016. White arrows indicate BP-FISH signals. Scale bar, 15μm. Histograms of proportion of extrachromosomal BP-FISH signal-positive cells on HF3016, HF3177 and PC3 cells (right panel). *p* values were determined by Mann-Whitney U test. (n > 100 cells per condition). The result is representative of an average of > 50 cells per sample studied from two separate experiments. **C.** Representative images of Casilio-labeled ecDNA(left panel). Scale bar, 10 μm. Histograms of targeting efficiency of Casilio-based labeling tool (right panel). *p* values were determined by Mann-Whitney U test. (n > 250 Casilio-transfected cells per condition). The result is representative of an average of > 50 cells per sample studied from four separate experiments. **D.** Representative two-color images of BP-FISH signal (Red) and Casilio-labeled ecDNA signal (green) (left panel). Scale bar, 10μm. Histograms of co-localized spots (right panel). (n > 20 cells per condition). **E.** ecDNA signal count per cell and MADs (left panel). *p* values indicating the homogeneity of variances between two groups (ecDNAs vs controls) were determined by Fligner-Killeen test. Individual *p* values of Fligner-Killeen test between each ecDNAs with Chr7 indicated in violin plot (right panel). The result is representative of an average of > 20 cells per sample.

To validate that the target breakpoints were specific to the ecDNA and were extrachromosomal, we used customized *in situ* probe sets targeting the compound sequence consisting of 10 nucleotides upstream and downstream of each breakpoint junction, enabling visual detection of breakpoints in metaphase cells^24^ (Methods and Supplementary Fig. 3A)^25^. A generic BAC library probe targeting the proximal region of the breakpoint on the same ecDNA was employed to distinguish ecDNA signal from the non-specific binding effect of probes. We also included a second neurosphere line, HF3177, which was derived from the recurrent glioblastoma from the same patient from whom HF3016 was established, therefore both cell lines were likely to share the same ecDNA amplifications. We have previously found that the PC3 prostate cancer cell line contains ecDNAs but very different in sequence from those in HF3016/HF3177, and therefore used PC3 as a negative control. The breakpoint-specific FISH (BP-FISH) analysis showed breakpoints co-labeled with two-color probes outside of chromosomes (Fig. 2B and Supplementary Fig. 3B), as well as breakpoint-negative extrachromosomal elements labeled with a BAC library probe suggesting additional ecDNA existed within each metaphase cell. The breakpoints were shared between primary (HF3016) and recurrent (HF3177) cell lines with different ratios (Fig. 2B, right panel). We observed some signal in the PC3 cells which is likely due to the relatively short length of the FISH probes, allowing for non-specific binding. The variability of breakpoint quantities was confirmed by BP-PCR (Supplementary Fig. 3C). These results showed that ecDNA-specific breakpoints can be leveraged to visualize ecDNA in single cells through fluorescence microscopy.

To engineer a live-cell ecDNA tracking system, we cloned breakpoint-specific sgRNAs with 25 PUFBS repeats. Co-transfection of sgRNAs, catalytically inactivated dCas9, and Clover-PUF fusion protein-expressing plasmids allows the enrichment of fluorescent signals at the targeted ecDNA breakpoint loci (Fig. 2A). To validate the targeting efficiency of breakpoint-specific sgRNAs, we performed an *in vitro* cleavage assay on HF3016 and HF3177 cells and confirmed on-target efficiency of sgRNA (Supplementary Fig. 4A).

To verify the labeling efficiency of the ecDNA tracking system, both HF3016 and HF3177 cell lines were co-transfected with three components: 1. breakpoint-specific sgRNA, 2. dCas9 and 3. Clover-PUF fusion protein expressing plasmid. Each component was prepared as an individual plasmid. Breakpoint-positive HF3016 and HF3177 cells, but not control PC3 cells, showed an abundance of nuclear spot signals, reflecting fluorescently labeled ecDNA breakpoints (Fig. 2C). Spot signals did not result from fluorescent molecules specifically aggregating in the nucleolus (Supplementary Fig. 4B). The mean targeting efficiency of the percentage of cells with fluorescent spots in HF3016 and HF3177 was 26% and 20% respectively, compared to an off-target 3% in PC3. Two-color imaging of BP-FISH and Casilio ecDNA labeling demonstrated that the spot signals derived from the Casilio-labeling method accurately mapped the specific ecDNA breakpoint (Fig. 2D and Supplementary Fig. 4C). Next, we applied this tool to evaluate the copy number distribution of the four different ecDNA breakpoints (ecEGFRx1, ecEGFR, ecCCAT1 and ecCCDC26). We included sgRNAs labeling an intronic region on chromosome 7 (Chr7) and the chr 3q29 gene *MUC4* as representative linear DNA controls. This confirmed the uneven ecDNA distribution with MADs of 1.5 to 3 in comparison to 1.5 and 1.5 for Chr7 and *MUC4,* respectively (Fig. 2E). While MADs of ecDNAs and controls were comparable, the copy number distribution of ecDNAs was significantly more variable than the Chr7 and MUC4 controls (Fig. 2E, right panel, p value = 0.0005, Fligner-Killeen homogeneity of variances test of ecDNAs vs controls).

### Spatiotemporal tracking of ecDNA shows uneven segregation of ecDNA during mitosis

Centromeres provide attachment sites for spindle microtubules to enable chromosome segregation and the acentromeric character of ecDNA therefore implies unequal segregation in mitosis^5, 26^. Using our Casilio - based ecDNA tracing system, we sought to evaluate the distribution of ecDNA following cell division. We monitored mitosis in HF3016 neurosphere cells with fluorescent labels attached via Casilio to ecDNAs. We found that the fluorescent signal faded during cytoplasmic division, which may be explained by the level of DNA compaction during metaphase^27^ or potential ecDNA clustering during mitosis ^28^, resulting in an inability of sgRNAs to bind target sequences (Fig. 3A). Once the telophase finished and the two daughter cells entered interphase, the fluorescent signals re-established again visualizing ecDNA molecules. We found that daughter cells often inherit different numbers of ecDNAs (Fig. 3A and Movie 1). We quantified fluorescent signals in offspring cells and observed that Chr7 and *MUC4* derived signals showed uniform segregation, reflected by a Pearson correlation of 1 or near 1, whereas the same analysis of ecDNA inheritance showed a marginal and non-significant correlation (Fig. 3B). This result demonstrates, unequivocally, that ecDNA segregates unevenly.

**Fig. 3.**
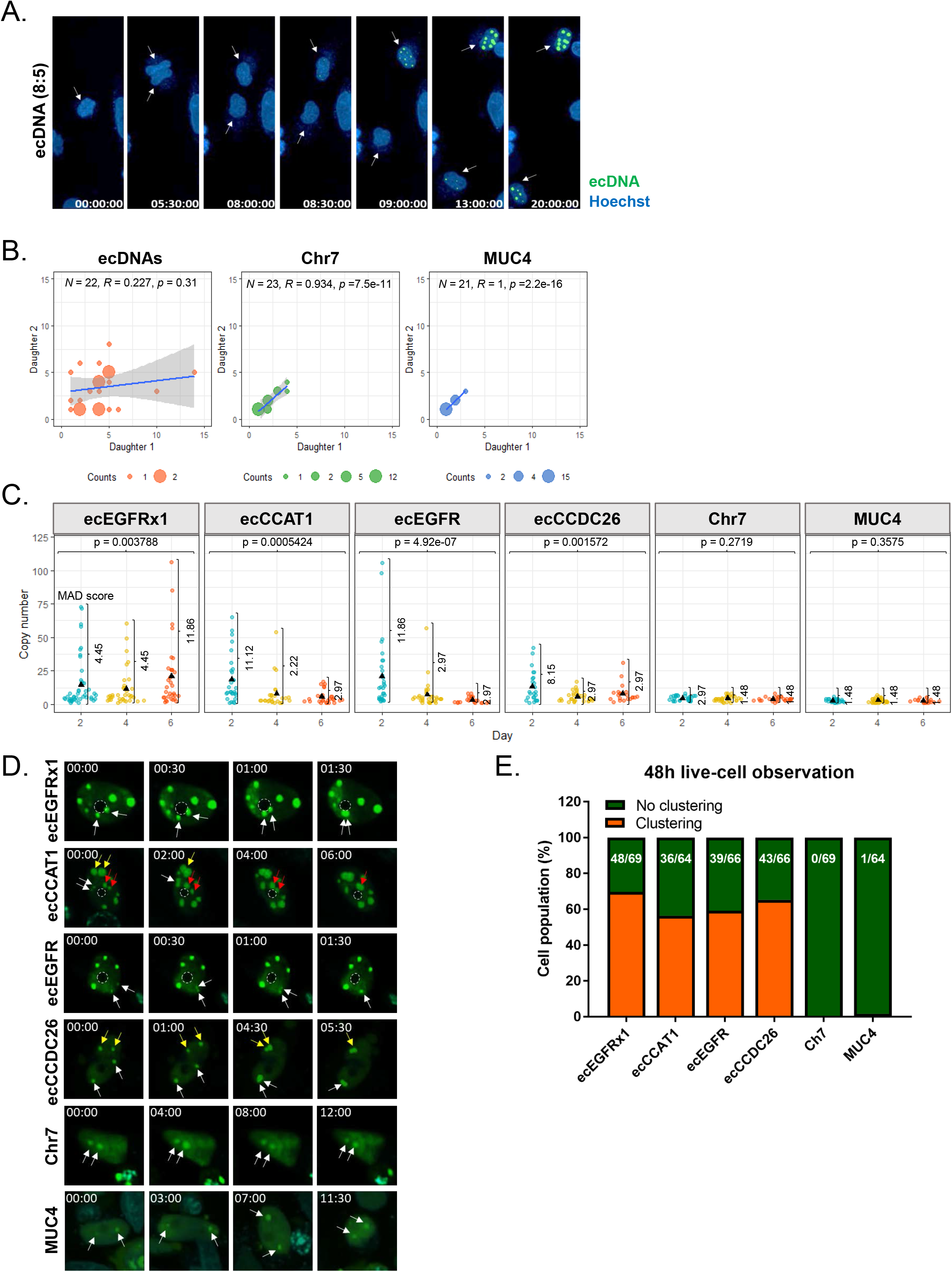
Spatiotemporal tracking of ecDNA shows uneven segregation of ecDNA during mitosis. **A.** Captured time-lapse images of ecDNA segregation during mitosis. Threshold-adjusted images of two daughter cells (D1 and D2) in Chr7 are displayed under the corresponding image. **B.** Copy number of ecDNAs, Chr7, and *MUC4* segregated into two daughter cells. (n > 20 dividing cells per each condition). Randomness of ecDNA segregation was determined by Pearson’s correlation test and the *p* value higher than 0.05 indicates the random distribution. **C.** Copy number distribution of ecDNAs, Chr7, and *MUC4* on three different days. Individual dot represents copy number count of a single-cell. The MADs are indicated. *p* values indicating the dynamic change of copy number variance between the days was determined by Fligner-Killeen test. The result is representative of a distribution of > 20 cells per sample. **D.** Captured time-lapse images of ecDNA clustering. The pair of arrows with the same color on each group showed the process of ecDNA clustering. (00:00 = Hour:Minute) **E.** The cell population with or without the clustering event of each Casilio signal was counted from the 48 h live-cell imaging data. The number of cells with the clustering event and total number of observed cells are indicated on each bar.

To obtain better insight into the discordant inheritance pattern of ecDNA, we determined the distribution of single-cell ecDNA and linear DNA copy numbers every two days, relative to the two day doubling time of HF3016 cells (Supplementary Fig. 5A). EcDNA copy number continued to be highly variable over several cell doublings, in comparison to the Chr7 and *MUC4* controls (Fig. 3C). In contrast, the average copy number of controls, Chr7 and *MUC4* – linear DNA amplicons on canonical chromosomes - remained stable, indicating that the DNA replication of these regions is tightly regulated by cell cycle^7, 16^. This result suggests that unlike linear DNA amplicons, ecDNA frequencies continuously fluctuate over time, and emphasizes the rapid mode of tumor evolution that ecDNA elements are able to direct in comparison to linear amplifications^8^.

The live-cell ecDNA tracking experiments provided a spatiotemporally dynamic feature of ecDNA within a single cell, but also suggested that ecDNA showed a propensity to physically cluster together, not involving the nucleolus (Fig. 3D). To confirm whether the spot signals are the hub of multiple ecDNA elements, we employed a dual-color labeling system (Supplementary Fig. 5B). Two sgRNAs were designed mapping to the same breakpoint to visualize ecDNA aggregation using green and red fluorescent molecules. We observed complete merging of yellow signals or closely assembled two colors indicating ecDNA clustering (Supplementary Fig. 5C). Moreover, ecDNA clustering was observed to take place in over 50% of cells within 48 hours of live cell imaging (Fig. 3E), expanding previous observations in anaphase cells^28^ and suggesting functional relevance.

### EcDNA clustering associates with RNA polymerase II activity

Recent studies have shown that ecDNA drives high levels of oncogene expression ^9, 29, 30^. We hypothesized that the generation of ecDNA clusters enhances transcriptional activity. First, we examined whether ecDNA clusters were more strongly associated with transcriptional activity through colocalization of nuclear bodies, including Cajal bodies and promyelocytic leukemia protein (PML) nuclear bodies ^31, 32^. These nuclear bodies are membrane-less subnuclear organelles and molecularly discrete entities that expedite specific nuclear functions by concentrating enzymes, substrates, and molecular machineries^33, 34^. Casilio-transfected cells were stained using Cajal/PML marker protein antibodies and a secondary antibody conjugated with red fluorescent molecule was used to capture the primary marker protein antibody. We compared colocalization between nuclear bodies and ecDNA or *MUC4* with Chr7 colocalization as a control. We observed a significantly higher Pearson’s coefficient value of colocalization for one of four ecDNA breakpoints (0.03, ecCCDC26 in Cajal body; 0.04, ecCCAT1 in PML body) in comparison to Chr7 (0.0, Cajal body; 0.0. PML body)(Supplementary Fig. 6A and Supplementary Fig. 7A), with only some ecDNAs colocalizing with nuclear bodies in cells with abundant presence of ecDNA (Fig. 4Ai and Bi). However, the fraction of cells containing ecDNAs colocalizing with nuclear bodies was substantially higher than the fraction of cells containing MUC4 or Chr7 colocalizing with nuclear bodies (Cajal bodies: 45%, 6% and 0% for ecDNAs, *MUC4,* and Chr7, respectively; PML bodies, 67%, 10% and 17% for ecDNAs, *MUC4*, and Chr7, respectively)(Fig. 4Aii and Bii). Since the amount of fluorescent signal is linearly correlated to the number of target copies and the amount of ecDNAs is much greater than the Chr7/*MUC4* control, we normalized by Casilio locus signal. This step did not change the result (0.07 ~ 0.24 in Cajal body and 0.09 ~ 0.16 in PML body, Mann-Whitney test, p-value < 3e-03 in all samples, ecDNAs vs Chr7; 0.03 in Cajal body and 0.05 in PML body, Mann-Whitney test, not significant, *MUC4* vs Chr7)(Fig. 4Aiii and Biii). This analysis also showed that a significantly higher proportion of ecDNA signal is merged with nuclear bodies compared with Chr7 (0.02 ~ 0.06 in Cajal body and 0.03 ~ 0.06 in PML body, Mann-Whitney test, p-value < 2e-03 in all samples, ecDNAs vs Chr7) while the proportion of *MUC4* area merged with nuclear bodies (0.01 in Cajal body and PML body, Mann-Whitney test, ns, *MUC4* vs Chr7) showed no significant differences (Supplementary Fig. 6B and Supplementary Fig. 7B).

**Fig. 4.**
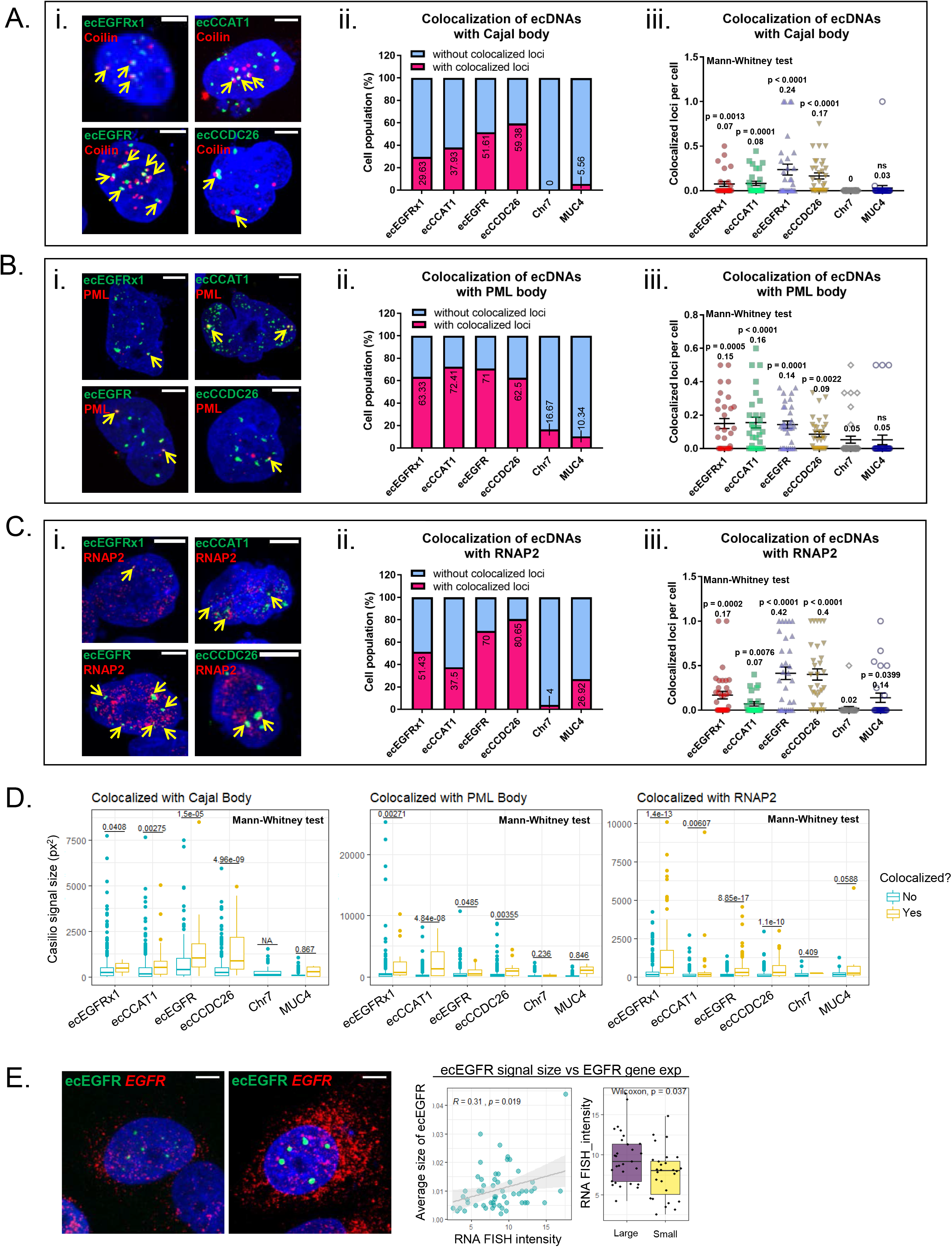
EcDNA bodies enhances transcriptional activity by recruiting RNA polymerase II (RNAPII). **A.** Representative images of Cajal body immunofluorescent staining, scale bar, 10 mm (i). Proportion of cells with or without the loci colocalized with Cajal body (ii). Colocalized loci with Cajal body per cell (iii). All value was normalized by each Casilio signal. The values of ecDNAs and *MUC4* were compared with Chr7. *p* values were determined by Mann-Whitney U test.. Average values are indicated under each *p* value. At least 28 single-cell images per group were analyzed. **B.** Representative images of PML body immunofluorescent staining, scale bar, 10 mm (i). Proportion of cells with or without the loci colocalized with PML body (ii). Colocalized loci with PML body per cell (iii). All value was normalized by each Casilio signal. The values of ecDNAs and *MUC4* were compared with Chr7. *p* values were determined by Mann-Whitney U test. Average values are indicated under each *p* value. At least 30 single-cell images per group were analyzed. **C.** Representative images of RNAPII immunofluorescent staining, scale bar, 10 mm (i). Proportion of cells with or without the loci colocalized with RNAPII (ii). Colocalized loci with RNAPII per cell (iii). All value was normalized by each Casilio signal. The values of ecDNAs and *MUC4* were compared with Chr7. *p* values were determined by Mann-Whitney U test. Average values are indicated under each *p* value. At least 25 single-cell images per group were analyzed. **D.** Comparison of ecDNA signal size by the matter of colocalization. The values of ecDNAs and *MUC4* were compared with Chr7. *p* values were determined by Mann-Whitney U test. The same images used in A-C were analyzed. **E.** Representative images of EGFR RNA FISH on Casilio-labeled cells with small ecDNA signals (left) and large ecDNA signals (right)), scale bar, 10 mm. Correlation between ecDNA signal size and EGFR gene expression (right panel). The scatter plot and Pearson’s correlation score showed a positive correlation. The bar plots represented the average EGFR gene expression in the cells with large size of ecEGFR signal and small size of ecEGFR signal. (median signal size = 0.009, large size ≥ 0.009, small size < 0.009). 49 single-cell images were analyzed.

Cajal and PML body have been reported to contain hyperphosphorylated RNA polymerase II (RNAPII), nominating them as sites of active mRNA transcription ^35, 36^. To determine whether ecDNA signal regions are actively transcribed, Casilio-transfected cells were stained with RNAPII antibody (Fig. 4Ci). Comparable to our observation on the colocalization with nuclear bodies, two of the four ecDNA breakpoint signals colocalized with RNAPII at higher frequency compared to the Chr7 control (Supplementary Fig. 8A). More than half of the cells (59.9 %) contained colocalizing RNAPII-ecDNAs (Fig. 4Cii). The number of colocalized loci per cell normalized by Casilio signal loci showed that the significantly higher number of ecDNA loci are colocalized with RNAPII compared with Chr7 (0.07 ~ 0.42, Mann-Whitney test, p-value < 8e-03 in all samples, ecDNAs vs Chr7). We also found significantly higher *MUC4*-RNAPII signal (0.14, Mann-Whitney test, p-value = 0.04, *MUC4* vs Chr7) in comparison to Chr7, reflecting the active transcription of a linear chromosomal gene (Fig. 4Ciii). The total Casilio signal area merged with RNAPII loci was measured and normalized by the area of each Casilio signal. We then defined the significantly higher proportion of ecDNA area as merged with RNAPII (0.01 ~ 0.08, Mann-Whitney test, p-value < 8e-03 in all samples, ecDNAs vs Chr7) compared with Chr7 (Supplementary Fig. 8B).

EcDNA clusters colocalizing with nuclear bodies or RNAPII were found to be significantly larger than the ecDNA without colocalization (Fig. 4D, Mann-Whitney test, p-value < 0.05 in all samples, ecDNAs vs Chr7). Chr7 and *MUC4* signals did not show significant differences in size, suggesting that the benefit of clustering to recruit functional transcriptional machinery is specific to ecDNA (Fig. 4D).

While we observed interactions between ecDNA and nuclear bodies at rates significantly higher than random (Supplementary Fig. 6A-B, Supplementary Fig. 7A-B), there was no significant linear correlation between the number of ecDNAs and the number of nuclear bodies, suggesting that ecDNA is not actively being trafficked towards the nuclear bodies (Supplementary Fig. 6C and Supplementary Fig. 7C). However, ecDNAs carrying *EGFR*-tagging breakpoints ecEGFRx1 or ecEGFR (Supplementary Fig. 2A-D), did show a positive correlation with RNAPII signals. We hypothesized that ecDNA-hubs form independent from other nuclear elements (Supplementary Fig. 8C), consisting of transcriptionally and RNAPII-bound ecDNA. As expected, we observed a positive correlation between *EGFR* FISH signal and EGFR protein expression using ImmunoFISH (Fig. 1E-F, lower panels). We sought to additionally examine *EGFR* RNA expression at the single-cell level. We observed that *EGFR* gene expression was positively correlated with the size of ecEGFR signals (Fig. 4E). We did not observe the same gene expression-Casilio signal size correlation for ecEGFRx1, which maps to ecDNAs containing only *EGFR* exon 1 (Supplementary Fig. 9B). Clustering of ecDNA may drive increased transcriptional activity of its cargo gene, with a spatial interaction advantage provided by clustering ecDNAs, thus exposing multiple ecDNA molecules to transcriptional machinery simultaneously.

## Discussion

Here, we take advantage of a CRISPR dead-Cas9 technique that enables ecDNA-specific fluorescent tagging to interrogate undiscovered ecDNA biology, including inheritance pattern and dynamic behavior. By doing so, we extend previous *in situ* single time point observations, which are limited in the level of advance they are able to provide. The CRISPR-based genome visualization shown here expands beyond single time point observations and to live-cell tracking, to demonstrate the longitudinal development of extrachromosomal DNA dynamics.

Previous approaches used three or more sgRNAs to map loci of interest ^11, 37^ which is not feasible when tagging a unique breakpoint sequence. The Casilio system^21^, which leverages the RNA binding domains of the PUF proteins fused with fluorescent molecule, enabled us to label a single non-repetitive target locus using a single sgRNA. Applying this technique for tracing ecDNA in the process of cell mitosis visualized unequal ecDNA segregation during cell division despite of the technical challenges of live-cell imaging, such as limited time frames (~48 hours) as a result of laser-induced cellular stress. The limited accessibility of DNA during mitosis where all genomic components including ecDNA are condensed, restricted the ability to visualize DNA from metaphase to telophase. Future technological developments are needed to overcome this limitation. The current results are based on transient transfection of effector molecules, including dCas9, guides and fluorescent tags. A CRISPR-based ecDNA labeling system in which the effectors are integrated into the genome and stably expressed in target cells will enable additional delineation of ecDNA behavior during tumor evolution. Despite these limitations, our results show that ecDNA, through random segregation during mitosis, enhances intratumoral diversity at the genomic level, and thus allowing ecDNA accumulation over the course of just a few cell cycles. This observation was reflected by fluctuations in ecDNA copy number distribution over subsequent cell cycles.

We observed that ecDNAs tend to converge in clusters leading to increased transcriptional activity. EcDNA hubs recruited RNAPII leading to active mRNA expression of cargo genes, highlighting an additional mechanism that explains the exceptionally high levels of ecDNA gene expression reported earlier^29, 30^. Our results compound recent discoveries of the wide-open chromatin accessibility of ecDNA^29^, the topological advantage of ecDNA for communicating with regulators^30^, and ecDNA-driven oncogenic genome remodeling^38^, and suggest that the advantage of tumor cells for maintaining ecDNA extends beyond simple dosage effects on cargo gene transcription. The three-dimensional topological orientation of genomic loci on linear chromosomes enables physical interaction between distal regulatory elements and gene promoters^39^. The circularization of oncogenes on ecDNAs increases enhancer interactions in ways restricted by insulators when on linear chromosomes^30^. The ecDNA clustering shown here creates additional interaction opportunities between ecDNA oncogene promoters and enhancers. Such ecDNA clusters can serve as transcriptional hotspots by sharing transcriptional machinery. Our results on ecDNA clusters reflect that the physical assembly of ecDNA may be required for optimal transcriptional activity. A comprehensive understanding of ecDNA clustering will help explain the biological roles of ecDNA contributing to gene expression, cell proliferation, and cell motility in cancer. Taken together, our observations build upon genome engineering technologies to provide new insights into ecDNA biology, a factor contributing to intratumoral heterogeneity. Defining mechanisms of ecDNA replication and assembly will be needed to understand how ecDNA can be leveraged for cancer therapeutics.

## Acknowledgements

This work was supported by the following NIH grants: R01 CA237208, R21 NS114873 and Cancer Center Support Grant P30 CA034196 (R.G.W.V); Department of Defense W81XWH1910246 (R.G.W.V); and grants from the Musella Foundation, the B*CURED Foundation and the Brain Tumour Charity (R.G.W.V). E.Y. is supported by a basic research fellowship from the American Brain Tumor Association (BRF1800014). K.C.J. is the recipient of an American Cancer Society Fellowship (130984-PF-17-141-01-DMC).

## Methods

### Human tumor specimens

Human glioma resection specimens (SMJL006, SMJL012, SMJL017 and SMJL018) were obtained from St. Michael’s Hospital. All tissue donations were approved by the Institutional Review Board of the Jackson Laboratory and clinical institutions involved. This work was performed in accordance with the Declaration of Helskinki principles.

### Cell cultures and cell lines

Patient-derived glioblastoma spheroids (HF3016 and HF3177) were cultured in neurophsere medium (NMGF): 500 ml DMEM/F12 medium (Invitrogen 11330) supplemented with N-2 (Gibco, 17502-048), 250 mg bovine serum albumin (BSA, Sigma, A4919), 12.5 mg gentamicin reagent (Gibco, 15710-064), 2.5 ml Antibiotic/Antimycotic (Invitrogen, 15240-062), 20 ng/ml EGF (Peprotech, 100-15), and 20 ng/ml bFGF (Peprotech, 100-18B). Human prostate cancer cell line PC3 was a gift from Dr. Paul Mischel at University of California at San Diego, and cultured in F12-K (ATCC, 30-2004) with 10% fetal bovine serum (FBS, VWR, 97068-085). All cultured cells were tested for Mycoplasma contamination before use with MycoAlert Mycoplasma Detection Kit (Lonza).

### FISH analysis

For patient tissues, the slightly thawed tissues were transferred to a positively charged glass slide by pressing against the surface of the specimen. The tissue slides were then immediately transferred into Carnoy’s fixative (3:1 methanol:glacial acetic acid, v/v), incubated at RT for 30 min and then air-dried. For interphase cell prep, neurospheres were dissociated into single cells and fixed in Carnoy’s fixative for 20 min, briefly washed with fixative and resuspended in fixative. Desired amounts of cells were then dropped onto the glass slide and air-dried. A hybridization buffer (Empire Genomics) mixed with EGFR-Chr7 probe (EGFR-CHR07-20-ORGR, Empire Genomics) was applied to the slides and the slides were denatured at 75°C for 5 min. The slides were then immediately transferred and incubated at 37°C overnight. The post-hybridization wash was with 0.4x SSC at 75°C for 3 min followed by a second wash with 2x SSC/0.05% Tween20 for 1 min. The slides were then briefly rinsed by water and air-dried. The VECTASHIELD mounting medium with DAPI (Vector Laboratories) was applied and the coverslip was mounted onto a glass slide. Tissue images were scanned under Leica STED 3X/DLS Confocal with 100x magnification. Z-stack acquired at 0.3-0.5 um step size was performed and all analysis conducted based on maximum intensity projection images of the 3D volume of the cells.

### Pan-cancer FISH analysis

We collected FISH images of four genes presented on ecDNA (MDM2, PDGFRA, EGFR, MYC) in six cell lines (CA718, GBM6, GBM39, HK359, MB411FH, and PC3) from GBM, medulloblastoma, and prostate cancer cell line (Supplementary Fig. 1, Red bar plots). We also collected FISH images of sixteen linearly amplified genes (TADA2A, CCND3, NFKBIA, RORC, EGFR, CCDN1, SBDS, HMGA2, BMP5, ARID5B, ERCC2, BRF2, IRF4, KCND3, TNFRSF13B, and IGFBP1) in sixteen cell lines (BP474, COLO205, DU145, EKVX, GSC11, H23, H322, HCC827, HCC1569, HK259, HOP62, OVCAR5, RPMI8226, SKBR3, SN12C, and SW620) from breast, colon, prostate, lung, GBM, ovary, hematopoietic, meduloblastoma, kidney cancer cell line (Supplementary Fig. 1, Blue bar plots). All images used here were obtained from https://figshare.com/s/6c3e2edc1ab299bb2fa0 and https://figshare.com/s/ab6a214738aa43833391.

### ImmunoFISH analysis

The patient tissue slides previously prepared by the tumor touch prep method were used. FISH analysis was performed on the slides as described above. After the last washing step with water, the slides were then incubated with blocking buffer (5% BSA/0.3% Triton X-100/1X PBS) at RT for 1 hr. Without washing step, EGFR antibody (#4267, Cell signaling, 1:100) diluted in antibody diluent (1% BSA/0.3% Triton X-100/1X PBS) was applied to the slide and incubated at RT for 1 hr. Then slides was washed with washing buffer (0.025% Tween-20/1X PBS) three times and the secondary antibody conjugated with Alexa 555 (ab150086, abcam, 1:1000) was applied. After 1 hr incubation, the slides were washed three times and counterstained with VECTASHIELD mounting medium with DAPI (Vector Laboratories). For neurospheres, 1.2 × 10^5^ cells were plated with 10% FBS-containing NMGF media into a glass-viewing area of confocal dish (VWR, 75856-740). The next day, cells were fixed with 4% PFA at RT for 10 min and briefly rinsed with 1X PBS. To allow permeabilization, fixed cells were incubated with washing buffer for 3 min. Then blocking buffer was applied and incubated at RT for 1 hr. Then the immunofluorescent staining process was performed as described above. After the final washing step, cells were fixed with 4% PFA for 10 min and dehydrated by incubating dishes in gradually increasing concentration (70%/80%/90%/100%) of EtOH. Air dried dishes were then used for performing FISH analysis as described above. Images were scanned under Leica STED 3X/DLS Confocal with 100x magnification. Z-stack acquired at 0.3-0.5 um step size was performed and all analysis conducted based on maximum intensity projection images of the 3D volume of the cells.

### Immunofluorescence staining

1.2 × 10^5^ cells of HF3016 were plated with 10% FBS-containing NMGF media into a glass-viewing area of confocal dish (VWR, 75856-740). The next day, cells were transfected with Casilio plasmids (83.3 ng of dCas9, 83.3 ng of sgRNA and 83.3 ng of Clover) with Lipofectamine 3000 (Invitrogen, L3000015). After 24 h, the media was changed to fresh media. After 48 h post-transfection, the dishes were briefly rinsed with 1X PBS three times and fixed with 4% PFA at RT for 15 min. The fixed cells were then rinsed with 1X PBS and permeabilized by incubating with washing buffer at RT for 5 min. The dishes were then incubated with blocking buffer ar RT for 1 hr and immediately the primary antibody (Coilin, ab87913; PML, ab96051; RNAPII, ab193468; Abcam) diluted in antibody diluent was applied and incubated at RT for 1 hr. The dishes were then washed with washing buffer three times and the secondary antibody conjugated with Alexa 555 was applied. After 1 hr incubation, the slides were washed three times and counterstained with VECTASHIELD mounting medium with DAPI. Images were scanned under Leica STED 3X/DLS Confocal with 100x magnification. Z-stack acquired at 0.3-0.5 um step size was performed and all analysis conducted based on maximum intensity projection images of the 3D volume of the cells.

### Breakpoint-specific PCR

Genomic DNA was isolated from each cell line using the QIAamp DNA mini kit (Qiagen 51304) or Quick DNA Mini Prep Plus kit (Zymo Research, D4068). Breakpoint-specific PCR was performed in an automated thermal cycler (BioRad, C1000 Touch Thermal Cycler). Each reaction mixture were prepared with AccuPrime Taq DNA Polymerase system (Invitrogen, 12339016). The PCR protocol was: denaturation at 94 °C for 5 min followed by 30 cycles comprising denaturation at 94 °C for 30 sec, primer annealing at 61 °C for 30 sec and DNA elongation at 68 °C for 1 min, further extension at 72 °C for 5 min and rapid cooling to 4 °C. PCR products were analyzed by 1% agarose gel electrophoresis and visualized under UV illumination after SYBR-Safe staining (Invitrogen, S33102). For Sanger sequencing, the PCR amplicons were extracted using Nucleospin Gel and PCR Clean-Up kit (Macherey-Nager, 740609). The bidirectional Sanger sequencing was performed by EtonBio.

### Breakpoint-specific FISH

Neurosphere cell cultures and PC3 were synchronized at metaphase by treating with 80 ng/ml Colcemid (Roche, 10-295-892-001) overnight. Cells were washed with PBS and incubated with 0.075 M KCl at 37 °C for 15 min. Samples were then fixed in Carnoy’s fixative (3:1 methanol:acetic acid, v/v) according to standard cytogenetic procedures. Metaphase cells were dropped onto glass slides. Breakpoint-specific FISH were performed using the ViewRNA ISH Cell Assay (Invitrogen, QVC001) for with custom-made breakpoint-specific probes targeting the junctions of extrachromosomal DNAs. All breakpoint-specific probes were designed to target the 10-nucleotides on each side of the breakpoint, spanning a total of 20 nucleotides. The regular FISH probes purified from BAC clones were purchased from Empire Genomics. We omitted the dehydration/rehydration step as well as the protease treatment step of the manufacturer’s protocol. To hybridize the probes to double-stranded extrachromosomal DNA, the probe mixture (breakpoint-specific probe and a related regular probe) in pre-warmed Probe Set Diluent QF buffer was applied to the metaphase slide and incubated at 82 °C for 5 min as the denaturation step. The serial hybridization was then performed following the manufacturer’s protocol. The images were captured on an Andor Dragonfly Spinning Disk system with iXon camera. An oil-immersion objective (60x) was used to observe metaphase chromosome bodies. As excitation laser, a 405 nm, a 488 nm, and a 561 nm were used to detect chromosome bodies, BAC probe, and breakpoint-specific probe respectively. Images were analyzed using Imaris image analysis software (Bitplane).

For colocalization FISH with EDTB, EDTB transfected cells were prepared as interphase in Carnoy’s fixative and the DNA denaturation time was reduced to 3 min to preserve the Clover signal. Images were captured on Opera Phenix High-Content Screening System (PerkinElmer). The water immersion objective (40x) was used. As excitation laser, a 405 nm, a 488 nm, and a 561 nm were used to detect nucleus, EDTB signal, and breakpoint-specific probe respectively. Images were analyzed using Harmony High-Content Imaging and Analysis Software (PerkinElmer).

### Cloning

The backbone vectors for cloning were gifted from Dr. Albert Cheng. The 5’-phosphorylated duplexed oligos for the guide sequences of each breakpoint were purchased from IDT. The sgRNA spacer sequences were digested via BbsI (NEB, R3539) and then were cloned into sgRNA-PUFBS expression vectors (single-color: pAC1373-pX-sgRNA-25xPUFBSa, dual-color: pCR8-sgRNA-15xPUFBSa and pCR8-sgRNA-15xPUFBSc). Guide sequences were then cloned into the digested vector using T4 ligase (Roche, 4898117001).

### Targeting efficiency test

Neurospheres were plated into 384-well plates at 8,000 cells per well with NMGF medium containing 2% FBS (5,000 cells per well for PC3). The next day, cells were transfected with 20 ng of dCas9, 20 ng of sgRNA and 20 ng of Clover with Lipofectamine 3000 (Invitrogen, L3000015). Cells were stained with CellMask Deep Red Plasma membrane stain (Invitrogen, C10046) and NucBlue Live ReadyProbe (Invitrogen, R37605) at 24 h post-transfection. Transfected cells were imaged on Opera Phenix High-Content Screening System (PerkinElmer). The water immersion objective (20x) was used. A 405 nm, a 488 nm and a 561 nm laser were used to detect nuclii, EDTB signal and plasma membrane respectively. Targeted cells by EDTB were detected and counted using the spot analysis algorithm of Harmony High-Content Imaging and Analysis Software (PerkinElmer).

### In vitro sgRNA test

The in vitro targeting efficiency of sgRNAs was tested using a Guide-it Complete sgRNA Screening System (Takara, 632636) according to the manufacturer’s instructions. Breakpoint-PCR amplicons were used as a targeting templates and incubated with appropriate sgRNA and recombinant Cas9 (rCas9) at 37°C for 1 hr. The cleaved fragments generated by sgRNA targeting were perceived on agarose gel.

### Live cell imaging

Neurospheres were plated into 100-mm tissue culture dishs at 1 million cells per dish with NMGF medium containing 10% FBS. The next day, cells were transfected with Casilio plasmids (1250 ng of dCas9, 1250 ng of sgRNA and 1250 ng Clover) with Lipofectamine 3000 (Invitrogen, L3000015). After 24 h, the cells were trypsinized and resuspended in DPBS including 10% FBS. To isolate EDTB-transfected cells, the GFP-positive cells were then sorted by FACSAria Fusion (BD Biosciences) with 130 μm nozzle. To recover the sorted cell fitness, the EDTB-transfected cells were then replated into 384-well plate at 5,000 cells per well with NMGF including 10% FBS and incubated at 37 °C overnight. The next day, cells were stained with CellMask Deep Red Plasma membrane stain (Invitrogen, C10046) and NucBlue Live ReadyProbe (Invitrogen, R37605). Live-cell imaging was performed on Opera Phenix High-Content Screening System (PerkinElmer) under 5% CO_2_ and 37 °C temperature. To avoid evaporation of medium, empty wells around the samples were filled with DPBS. All the images were acquired by taking 6 – 8 different focal planes, and shown as a maximum intensity projection. To track extrachromosomal DNA dynamics, images were acquired every 30 min for 24 h or 48 h with 20x magnification. Time-lapse movies and snapshot images were generated and analyzed by Harmony High-Content Imaging and Analysis Software (PerkinElmer).

### Dual-color ecDNA labeling experiment

3 × 10^5^ cells of HF3016 were plated with 10% FBS-containing NMGF media into a glass-viewing area of confocal dish (VWR, 75856-742). The next day, cells were transfected with Casilio plasmids (400ng of dCas9, 200 ng of sgRNA-PUFBSa, 200 ng of sgRNA-PUFBSc, 200 ng Clover-PUFa and 200 ng mRuby-PUFc) with Lipofectamine 3000 (Invitrogen, L3000015). After 24 h, the media was changed to fresh media. After 48 h post-transfection, the dishes were briefly rinsed with 1X PBS three times and fixed with 4% PFA at RT for 15 min. The fixed cells were then rinsed with 1X PBS and counterstained with VECTASHIELD mounting medium with DAPI. Images were scanned under Leica STED 3X/DLS Confocal with 100x magnification. Z-stack acquired at 0.3-0.5 um step size was performed and all analysis conducted based on maximum intensity projection images of the 3D volume of the cells.

### Neurosphere doubling time test

HF3016 cells were plated in individual wells of a 96-well plate and viable cells were quantified using CellTiter-Glo 3D Cell Viability Assay (Promega) in triplicate wells as per manufacturer’s instructions. Luminescence readings, which represented viable cells, were taken on a Cytation 3 Cell Imaging Multi-Mode Reader (BioTek).

### Test for the dynamic change of ecDNA copy number distribution

To image Casilio-transfected cells every two days (Day2, Day4, and Day6), three sets of cells were prepared. 1.2 × 10^5^ cells of HF3016 were plated with 10% FBS-containing NMGF media into a glass-viewing area of confocal dish (VWR, 75856-740). The next day, cells were transfected with Casilio plasmids (83.3 ng of dCas9, 83.3 ng of sgRNA and 83.3 ng of Clover) with Lipofectamine 3000 (Invitrogen, L3000015). After 24 h, the media was changed to fresh media. After two days post-transfection, the dishes of the first set were briefly rinsed with 1X PBS three times and fixed with 4% PFA at RT for 15 min. The fixed cells were then rinsed with 1X PBS and counterstained with VECTASHIELD mounting medium with DAPI. Images were scanned under Leica STED 3X/DLS Confocal with 100x magnification. Z-stack acquired at 0.3-0.5 um step size was performed and all analysis conducted based on maximum intensity projection images of the 3D volume of the cells. The second set of cells were processed after four days post-transfection and the third set of cells were processed after six days post-transfection.

### EGFR RNA FISH

Fluorescence (Quasar 670 Dye)-conjugated EGFR DesignReady probe was purchased from Biosearch Technologies. RNA FISH was performed on HF3016 cells transfected with Casilio-labeling components using manufacturer’s protocol. Images were scanned under Leica STED 3X/DLS Confocal with 100x magnification. Z-stack acquired at 0.3-0.5 um step size was performed and all analysis conducted based on maximum intensity projection images of the 3D volume of the cells.

### Image analysis and data availability

Macro scripting of FIJI was used for automated analysis. Speckle inspector function in the BioVoxxel plugin was used for counting copy number of fluorescent signals. Colocalization analysis was done using the JACoP plugin (Pearson’s correlation test), the Image Calulator, and the Analyze Particle function of FIJI.

### Statistics (GraphPad)

All sample sizes and statistical methods were indicated in the corresponding figure or figure legends. All data was tested for normality using the D’Agostino-Pearson omnibus test. According to the results of the normality test, all data in this study that was not normally distributed wasthen run through the Mann-Whitney U test (for two groups). The homogeneity of variances between groups was determined by Levene’s test or Fligner-Kileen test. All statistical tests are two-sided. All plots are shown with median, upper and lower quartiles. All statistical test were performed in GraphPad Prism 7 or R version 4.0.2.

---

**Supplementary Fig. 1 | Comparison of copy number distribution between ecDNA genes and linearly amplified genes. A.** Individual copy number distribution on various types of cancer cell lines. Circularly amplified genes are considered as ecDNA gene and indicated by red. Linearly amplified genes were indicated by blue. The MADs are indicated at the top of each histogram.

**Supplementary Fig. 2 | The validation of breakpoint junction sequences. A-D.** ecDNA structures anticipated by AmpliconArchitect. Each ecDNA is named based on its cargo gene (A. ecEGFRx1 (exon 1); B. ecCCAT1; C. ecEGFR; D. ecCCDC26). **E.** Gel-images of BP-PCR across breakpoint junctions. **F.** Chromatograms of Sanger sequencing results for each breakpoint. The target specificity score was determined by CRISPOR.

**Supplementary Fig. 3 | The validation of breakpoint sequences. A.** Schematic illustration of BP-FISH method. **B.** Comprehensive single-channel images of BP-FISH results. Scale bar, 15 μm. C. Gel-images of comparative BP-PCR performed on HF3016 and HF3177 (left panel). Input genomic DNAs were quantified by quantitative PCR (qPCR) on GAPDH (right panel).

**Supplementary Fig. 4 | Specificity test of breakpoint-targeting sgRNAs. A.** Schematic illustration of the workflows of the specificity test (upper panel). Gel-images of BP-PCR amplicons sufficiently targeted by sgRNAs (lower panel). **B.** Comprehensive single-channel images of Casilio-labeled cells.The nucleolus is indicated by red arraows. Scale bar, 20μm. **C.** Comprehensive single-channel images of co-labeling of ecDNA with two colors (red = BP-FISH, green = Casilio). Scale bar, 10μm.

**Supplementary Fig. 5 | HF3016 doubling time test and dual-color ecDNA labeling system. A.** The doubling time (DT) of HF3016 cells were determined by cell viability assay. **B.** Schematic illustration of dual-color ecDNA labeling system. **C.** Representative dual-color labeling experiments. Scale bar, 10μm.

**Supplementary Fig. 6 | Colocalization of ecDNAs with Cajal body. A.** Average pearson’s correlation score between ecDNA signal loci and Cajal body loci. The values of ecDNAs and *MUC4* were compared with Chr7. *p* values were determined by Mann-Whitney U test **B.** Casilio signal area merged with Cajal body marker was normalized by each Casilio signal area. The values of ecDNAs and *MUC4* were compared with Chr7. *p* values were determined by Mann-Whitney U test. Average values are indicated under each *p* value. **C.** Correlation between copy number of ecDNAs and Cajal body count. Correlation score and *p* values were determined by Pearson’s correlation test. At least 28 single-cell images per group were analyzed.

**Supplementary Fig. 7 | Colocalization of ecDNAs with PML body. A.** Average pearson’s correlation score between ecDNA signal loci and PML body loci. The values of ecDNAs and *MUC4* were compared with Chr7. *p* values were determined by Mann-Whitney U test. **B.** Casilio signal area merged with PML body marker was normalized by each Casilio signal area. The values of ecDNAs and *MUC4* were compared with Chr7. *p* values were determined by Mann-Whitney U test. Average values are indicated under each *p* value. **C.** Correlation between copy number of ecDNA and PML body count. Correlation score and *p* values were determined by Pearson’s correlation test. At least 30 single-cell images per group were analyzed.

**Supplementary Fig. 8 | Colocalization of ecDNAs with RNAPII. A.** Average pearson’s correlation score between ecDNA signal loci and RNAPII loci. The values of ecDNAs and *MUC4* were compared with Chr7. *p* values were determined by Mann-Whitney U test. **B.** Casilio signal area merged with RNAPII marker was normalized by each Casilio signal area. The values of ecDNAs and *MUC4* were compared with Chr7. *p* values were determined by Mann-Whitney U test. Average values are indicated under each *p* value. **C.** Correlation between copy number of ecDNA and RNAPII count. Correlation score and *p* values were determined by Pearson’s correlation test. The positively correlated cases are marked with red star. At least 25 single-cell images per group were analyzed.

**Supplementary Fig. 9 | Correlation between ecDNA clustering and EGFR gene expression. A.** Correlation between copy number of ecEGFRx1 signals and EGFR gene expression. **B-F.** Correlation between copy number of each Casilio signals and EGFR gene expression (left panels). Correlation between the average signal size of each Casilio signals and EGFR gene expression (right panels). The correlation was determined by Pearson’s correlation test. The bar plots represented the comparison of average EGFR gene expression by copy number and signal size (lower panels). The category was determined by its median value. At least 40 single-cell images per group were analyzed.

## References

1. Gillies, R.J., Verduzco, D. & Gatenby, R.A. Evolutionary dynamics of carcinogenesis and why targeted therapy does not work. Nature Reviews Cancer 12, 487–493 (2012).

2. Andor, N. et al. Pan-cancer analysis of the extent and consequences of intratumor heterogeneity. Nature medicine 22, 105 (2016).

3. Amirouchene-Angelozzi, N., Swanton, C. & Bardelli, A. Tumor Evolution as a Therapeutic Target. Cancer discovery (2017).

4. Li, Y. et al. Patterns of somatic structural variation in human cancer genomes. Nature 578, 112–121 (2020).

5. Turner, K.M. et al. Extrachromosomal oncogene amplification drives tumour evolution and genetic heterogeneity. Nature 543, 122–125 (2017).

6. Nathanson, D.A. et al. Targeted therapy resistance mediated by dynamic regulation of extrachromosomal mutant EGFR DNA. Science 343, 72–76 (2014).

7. Verhaak, R.G.W., Bafna, V. & Mischel, P.S. Extrachromosomal oncogene amplification in tumour pathogenesis and evolution. Nature reviews. Cancer 19, 283–288 (2019).

8. Decarvalho, A.C. et al. Discordant inheritance of chromosomal and extrachromosomal DNA elements contributes to dynamic disease evolution in glioblastoma. Nature genetics 50, 708–717 (2018).

9. Kim, H. et al. Extrachromosomal DNA is associated with oncogene amplification and poor outcome across multiple cancers. Nature Genetics (2020).

10. Wahl, G.M. The Importance of Circular DNA in Mammalian Gene Amplification. Cancer Research 49, 1333–1340 (1989).

11. Chen, B. et al. Dynamic Imaging of Genomic Loci in Living Human Cells by an Optimized CRISPR/Cas System. Cell 155, 1479–1491 (2013).

12. Wang, H. et al. CRISPR-Mediated Programmable 3D Genome Positioning and Nuclear Organization. Cell 175, 1405–1417.e1414 (2018).

13. Wang, H. et al. CRISPR-mediated live imaging of genome editing and transcription. Science 365, 1301–1305 (2019).

14. Qin, P. et al. Live cell imaging of low-and non-repetitive chromosome loci using CRISPR-Cas9. Nature Communications 8, 14725 (2017).

15. Clow, P.A., Jillette, N., Zhu, J.J. & Cheng, A.W. CRISPR-mediated Multiplexed Live Cell Imaging of Nonrepetitive Genomic Loci. bioRxiv, 2020.2003.2003.974923 (2020).

16. Bailey, C., Shoura, M.J., Mischel, P.S. & Swanton, C. Extrachromosomal DNA—relieving heredity constraints, accelerating tumour evolution. Annals of Oncology 31, 884–893 (2020).

17. Johnson, K.C. et al. Single-cell multimodal glioma analyses reveal epigenetic regulators of cellular plasticity and environmental stress response. bioRxiv, 2020.2007.2022.215335 (2020).

18. Lundberg, G. et al. Binomial mitotic segregation of MYCN-carrying double minutes in neuroblastoma illustrates the role of randomness in oncogene amplification. PLoS One 3, e3099 (2008).

19. Patel, A.P. et al. Single-cell RNA-seq highlights intratumoral heterogeneity in primary glioblastoma. Science 344, 1396–1401 (2014).

20. Singh, D.K. et al. Patterns of basal signaling heterogeneity can distinguish cellular populations with different drug sensitivities. Molecular systems biology 6, 369 (2010).

21. Cheng, A.W. et al. Casilio: a versatile CRISPR-Cas9-Pumilio hybrid for gene regulation and genomic labeling. Cell Research 26, 254–257 (2016).

22. Deshpande, V. et al. Exploring the landscape of focal amplifications in cancer using AmpliconArchitect. Nature Communications 10, 392 (2019).

23. Concordet, J.-P. & Haeussler, M. CRISPOR: intuitive guide selection for CRISPR/Cas9 genome editing experiments and screens. Nucleic Acids Research 46, W242–W245 (2018).

24. Zhang, X. et al. In situ analysis of intrahepatic virological events in chronic hepatitis B virus infection. The Journal of clinical investigation 126, 1079–1092 (2016).

25. You, X. et al. Neural circular RNAs are derived from synaptic genes and regulated by development and plasticity. Nature neuroscience 18, 603–610 (2015).

26. Cleveland, D.W., Mao, Y. & Sullivan, K.F. Centromeres and kinetochores: from epigenetics to mitotic checkpoint signaling. Cell 112, 407–421 (2003).

27. Hsiung, C.C.S. et al. Genome accessibility is widely preserved and locally modulated during mitosis. Genome Res 25, 213–225 (2015).

28. Kanda, T., Sullivan, K.F. & Wahl, G.M. Histone-GFP fusion protein enables sensitive analysis of chromosome dynamics in living mammalian cells. Current biology : CB 8, 377–385 (1998).

29. Wu, S. et al. Circular ecDNA promotes accessible chromatin and high oncogene expression. Nature 575, 699–703 (2019).

30. Morton, A.R. et al. Functional Enhancers Shape Extrachromosomal Oncogene Amplifications. Cell 179, 1330–1341.e1313 (2019).

31. Wang, Q. et al. Cajal bodies are linked to genome conformation. Nature Communications 7, 10966 (2016).

32. Bernardi, R. & Pandolfi, P.P. Structure, dynamics and functions of promyelocytic leukaemia nuclear bodies. Nature Reviews Molecular Cell Biology 8, 1006–1016 (2007).

33. Mao, Y.S., Zhang, B. & Spector, D.L. Biogenesis and function of nuclear bodies. Trends Genet 27, 295–306 (2011).

34. Sleeman, J.E. & Trinkle-Mulcahy, L. Nuclear bodies: new insights into assembly/dynamics and disease relevance. Current Opinion in Cell Biology 28, 76–83 (2014).

35. Gall, J.G., Bellini, M., Wu, Z.a. & Murphy, C. Assembly of the Nuclear Transcription and Processing Machinery: Cajal Bodies (Coiled Bodies) and Transcriptosomes. Molecular Biology of the Cell 10, 4385–4402 (1999).

36. Kiesslich, A., von Mikecz, A. & Hemmerich, P. Cell cycle-dependent association of PML bodies with sites of active transcription in nuclei of mammalian cells. Journal of structural biology 140, 167–179 (2002).

37. Maass, P.G., Barutcu, A.R., Weiner, C.L. & Rinn, J.L. Inter-chromosomal Contact Properties in Live-Cell Imaging and in Hi-C. Mol Cell 69, 1039–1045 e1033 (2018).

38. Koche, R.P. et al. Extrachromosomal circular DNA drives oncogenic genome remodeling in neuroblastoma. Nature Genetics 52, 29–34 (2020).

39. Dekker, J. & Mirny, L. The 3D Genome as Moderator of Chromosomal Communication. Cell 164, 1110–1121 (2016).

