## Supplementary Figures for "Live-cell imaging shows uneven segregation of extrachromosomal DNA elements and transcriptionally active extrachromosomal DNA clusters in cancer"

### Supplementary Figure 1.

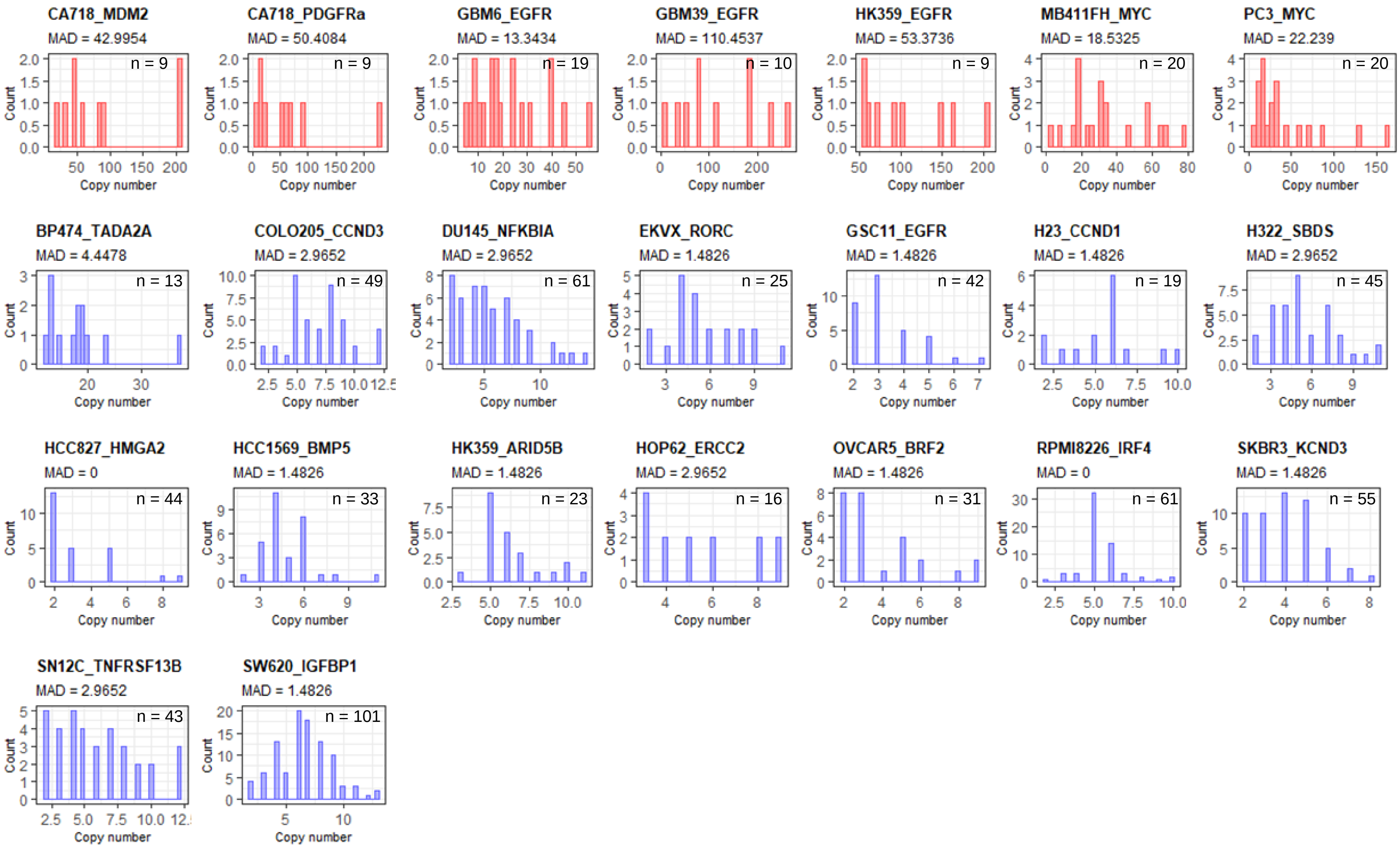


**Supplementary Fig. 1 | Comparison of copy number distribution between ecDNA genes and linearly amplified genes. A.** Individual copy number distribution on various types of cancer cell lines. Circularly amplified genes are considered as ecDNA gene and indicated by red. Linearly amplified genes were indicated by blue. The MADs are indicated at the top of each histogram.

### Supplementary Figure 2.

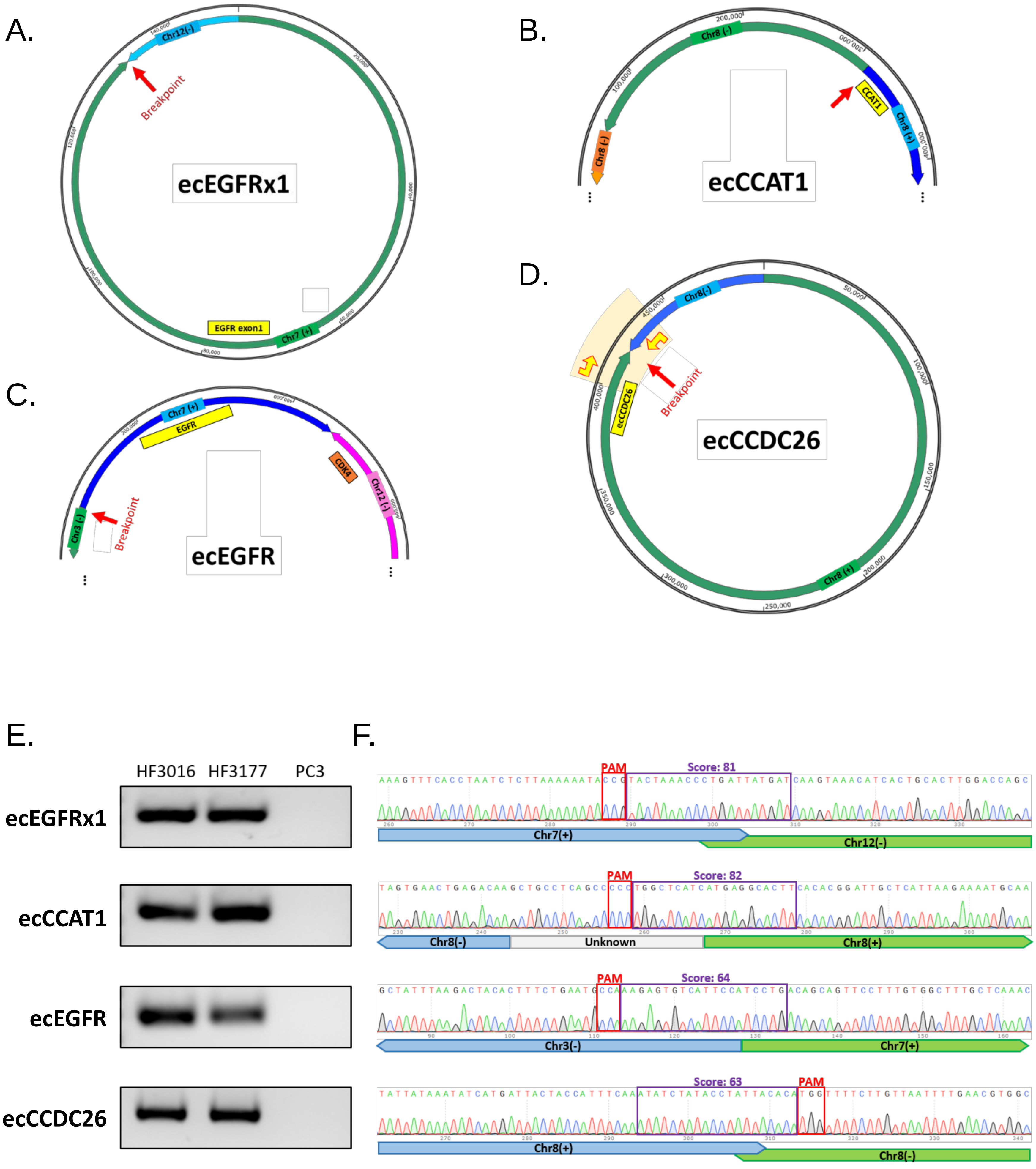


**Supplementary Fig. 2 | The validation of breakpoint junction sequences. A-D.** ecDNA structures anticipated by AmpliconArchitect. Each ecDNA is named based on its cargo gene (A. ecEGFRx1 (exon 1); B. ecCCAT1; C. ecEGFR; D. ecCCDC26). **E.** Gel-images of BP-PCR across breakpoint junctions. **F.** Chromatograms of Sanger sequencing results for each breakpoint. The target specificity score was determined by CRISPOR.

### Supplementary Figure 3.

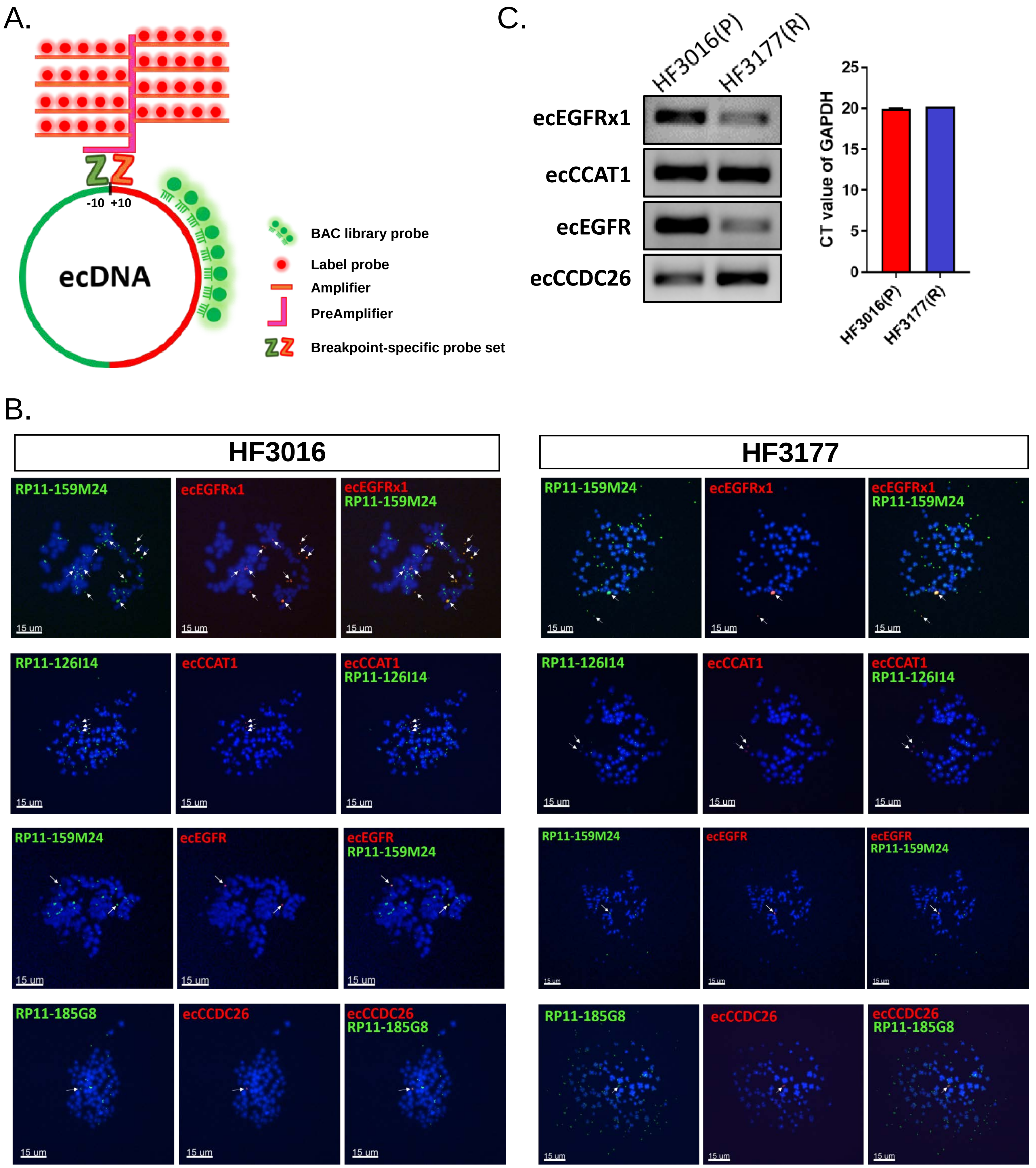


**Supplementary Fig. 3 | The validation of breakpoint sequences. A.** Schematic illustration of BP-FISH method. **B.** Comprehensive single-channel images of BP-FISH results. Scale bar, 15 μm. C. Gel-images of comparative BP-PCR performed on HF3016 and HF3177 (left panel). Input genomic DNAs were quantified by quantitative PCR (qPCR) on GAPDH (right panel).

### Supplementary Figure 4.

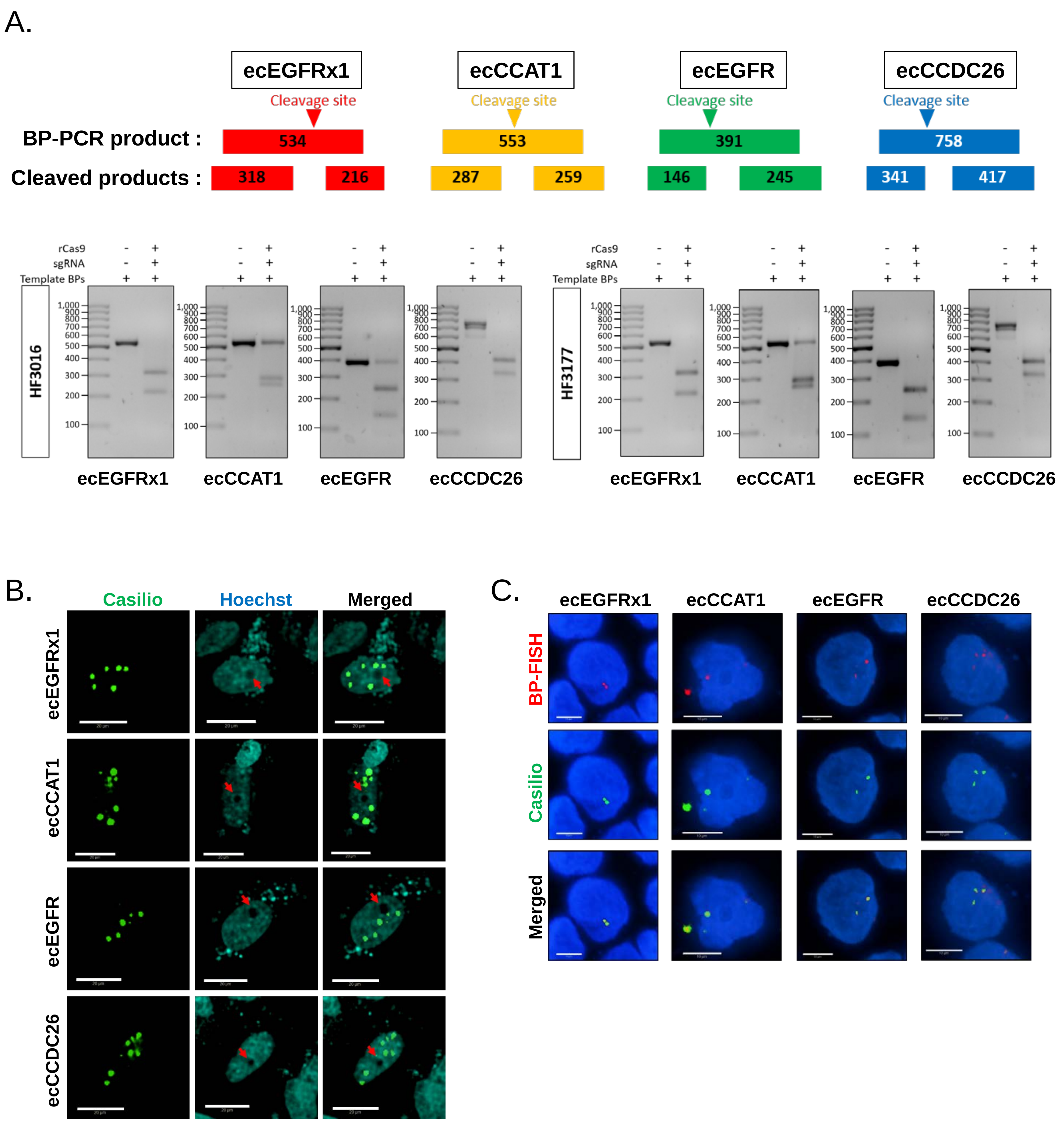


**Supplementary Fig. 4 | Specificity test of breakpoint-targeting sgRNAs. A.** Schematic illustration of the workflows of the specificity test (upper panel). Gel-images of BP-PCR amplicons sufficiently targeted by sgRNAs (lower panel). **B.** Comprehensive single-channel images of Casilio-labeled cells.The nucleolus is indicated by red arraows. Scale bar, 20µm. **C.** Comprehensive single-channel images of co-labeling of ecDNA with two colors (red = BP-FISH, green = Casilio). Scale bar, 10µm.

### Supplementary Figure 5.

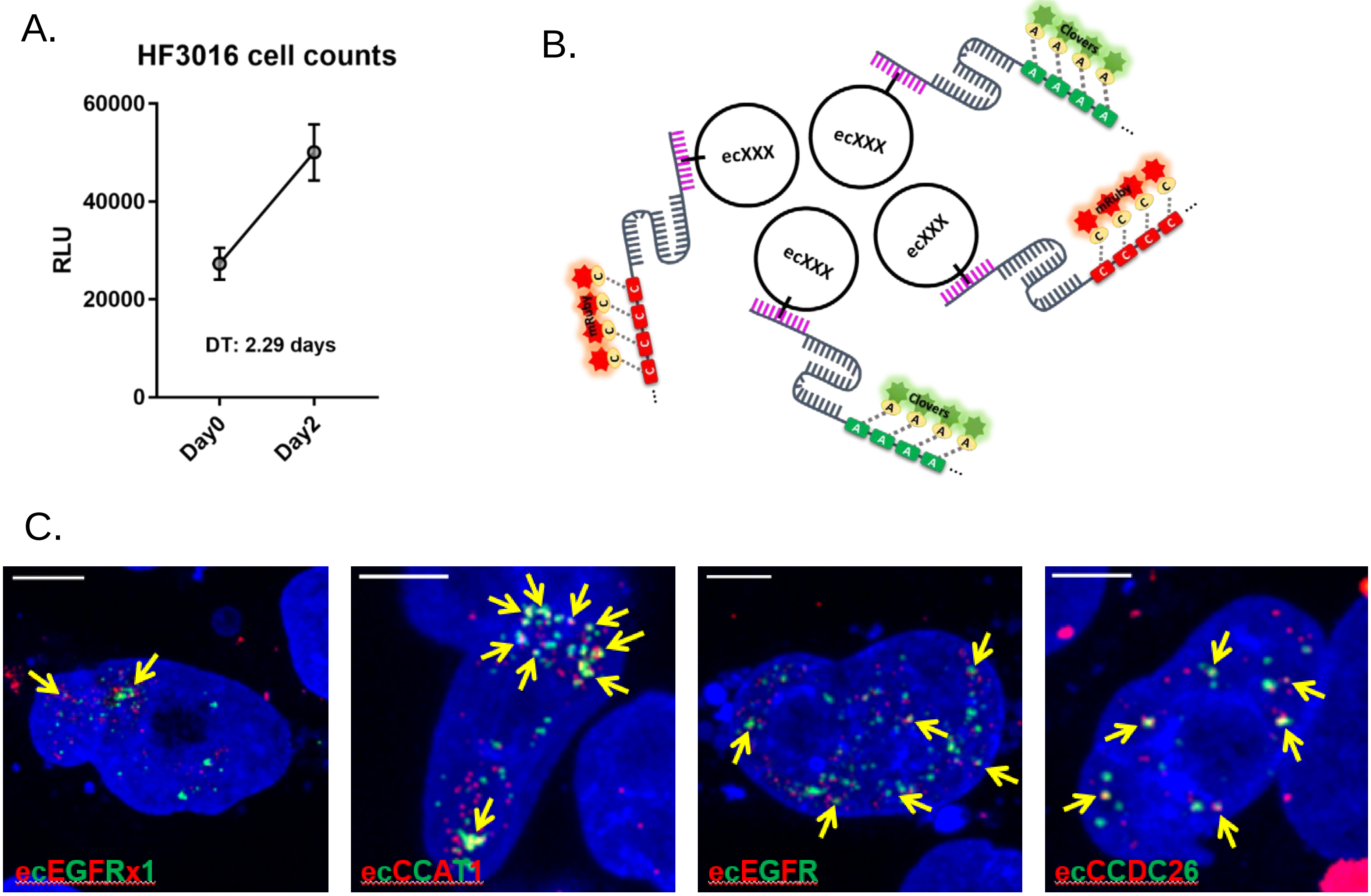


**Supplementary Fig. 5 | HF3016 doubling time test and dual-color ecDNA labeling system. A.** The doubling time (DT) of HF3016 cells were determined by cell viability assay. **B.** Schematic illustration of dual-color ecDNA labeling system. **C.** Representative dual-color labeling experiments. Yellow arrow indicates ecDNA clustering. Scale bar, 10μm.

### Supplementary Figure 6.

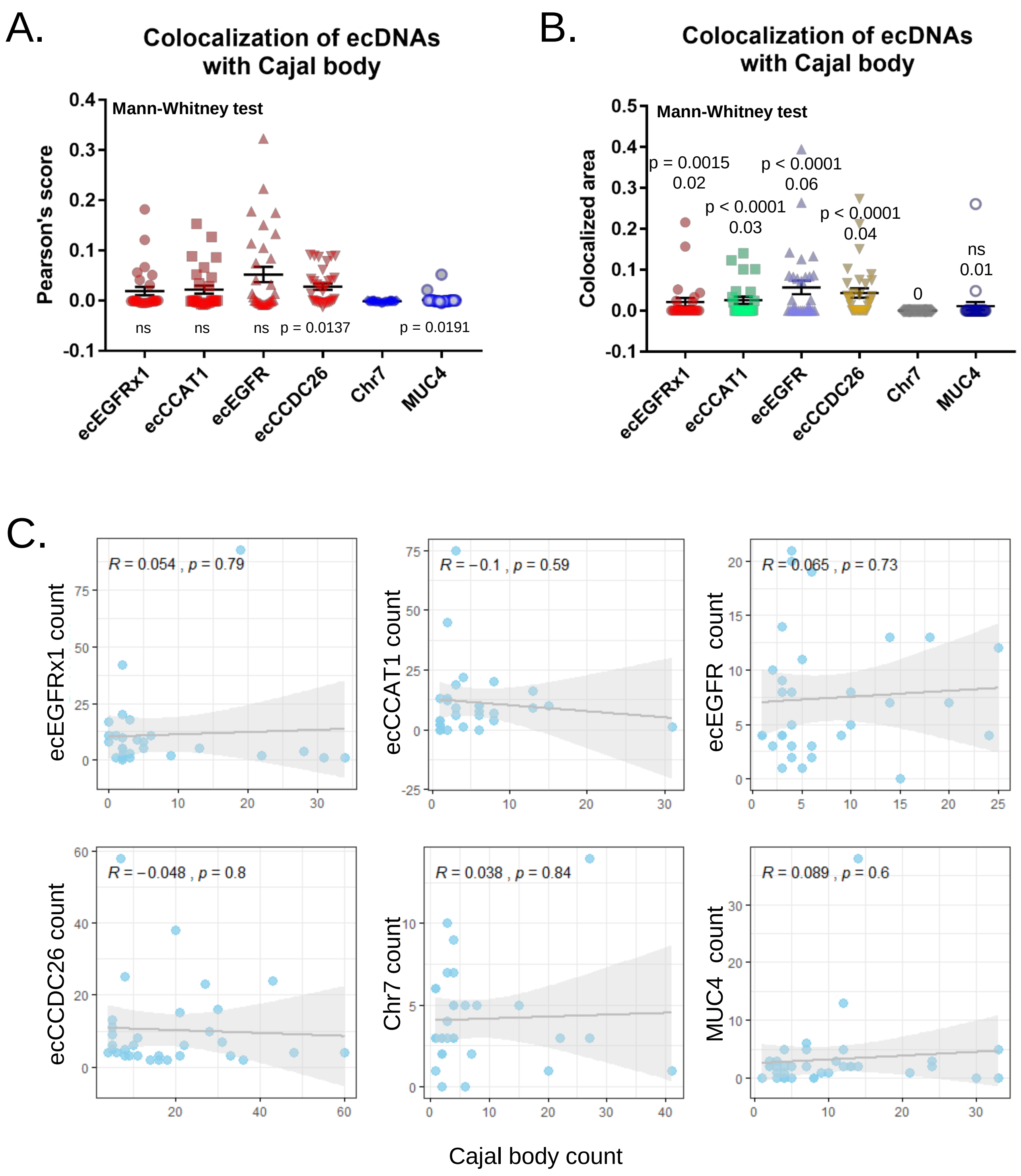


**Supplementary Fig. 6 | Colocalization of ecDNAs with Cajal body. A.** Average pearson's correlation score between ecDNA signal loci and Cajal body loci. The values of ecDNAs and *MUC4* were compared with Chr7. *p* values were determined by Mann-Whitney U test **B.** Casilio signal area merged with Cajal body marker was normalized by each Casilio signal area. The values of ecDNAs and *MUC4* were compared with Chr7. *p* values were determined by Mann-Whitney U test. Average values are indicated under each *p* value. **C.** Correlation between copy number of ecDNAs and Cajal body count. Correlation score and *p* values were determined by Pearson's correlation test. At least 28 single-cell images per group were analyzed.

### Supplementary Figure 7.

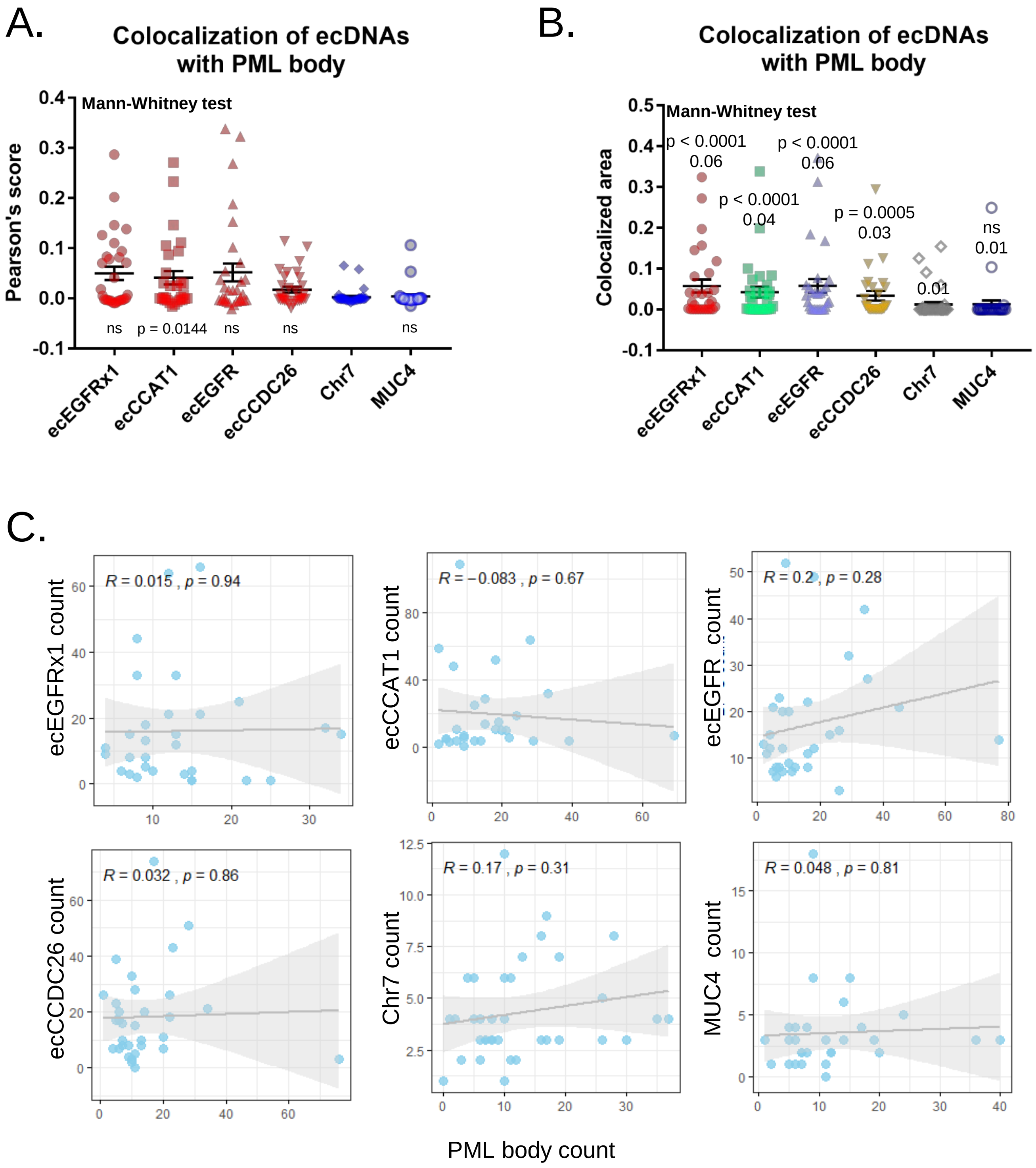


**Supplementary Fig. 7 | Colocalization of ecDNAs with PML body. A.** Average pearson's correlation score between ecDNA signal loci and PML body loci. The values of ecDNAs and *MUC4* were compared with Chr7. *p* values were determined by Mann-Whitney U test. **B.** Casilio signal area merged with PML body marker was normalized by each Casilio signal area. The values of ecDNAs and *MUC4* were compared with Chr7. *p* values were determined by Mann-Whitney U test. Average values are indicated under each *p* value. **C.** Correlation between copy number of ecDNA and PML body count. Correlation score and *p* values were determined by Pearson's correlation test. At least 30 single-cell images per group were analyzed.

### Supplementary Figure 8.

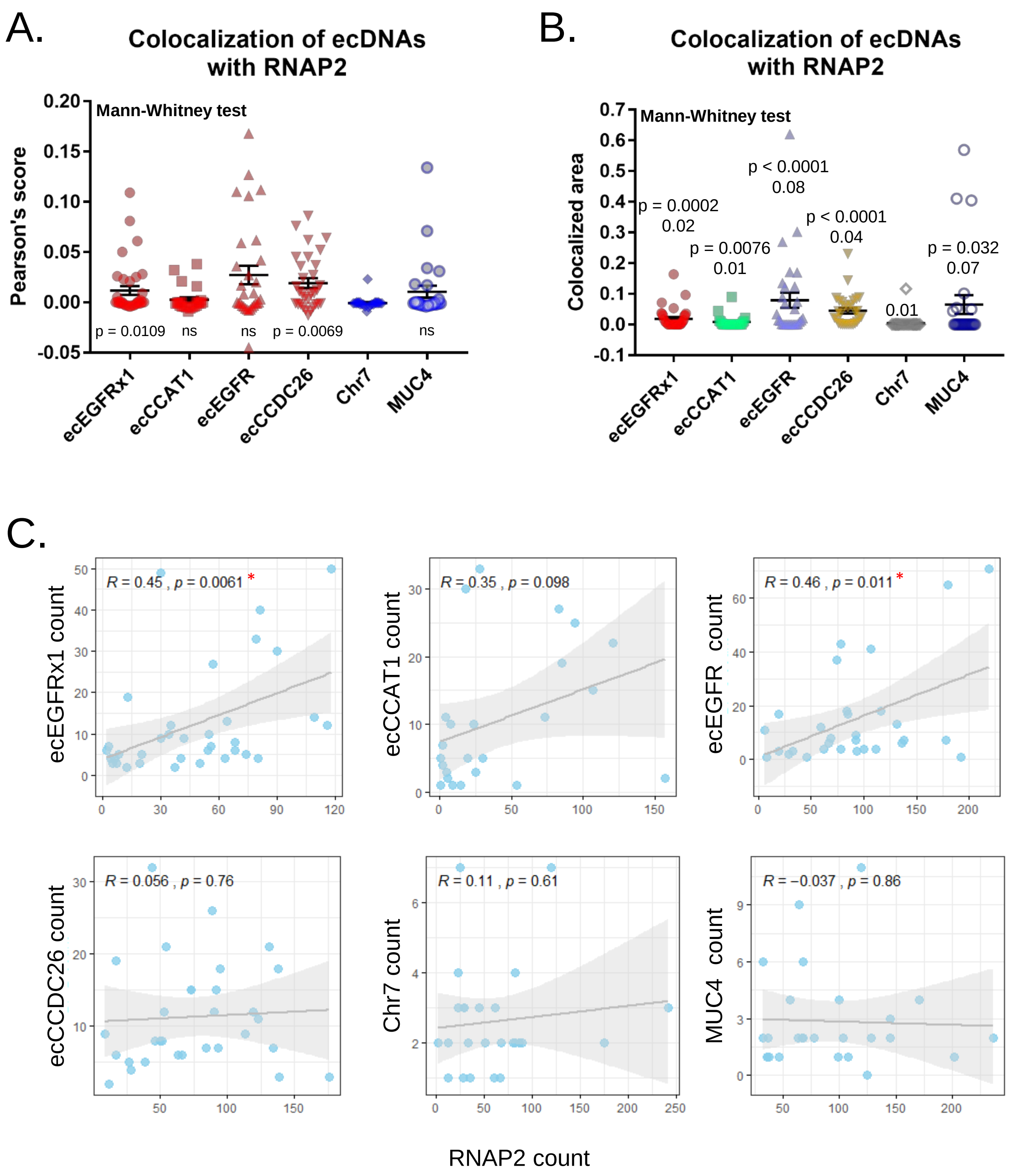


**Supplementary Fig. 8 | Colocalization of ecDNAs with RNAPII. A.** Average pearson's correlation score between ecDNA signal loci and RNAPII loci. The values of ecDNAs and *MUC4* were compared with Chr7. *p* values were determined by Mann-Whitney U test. **B.** Casilio signal area merged with RNAPII marker was normalized by each Casilio signal area. The values of ecDNAs and *MUC4* were compared with Chr7. *p* values were determined by Mann-Whitney U test. Average values are indicated under each *p* value. **C.** Correlation between copy number of ecDNA and RNAPII count. Correlation score and *p* values were determined by Pearson's correlation test. The positively correlated cases are marked with red star. At least 25 single-cell images per group were analyzed.

### Supplementary Figure 9.

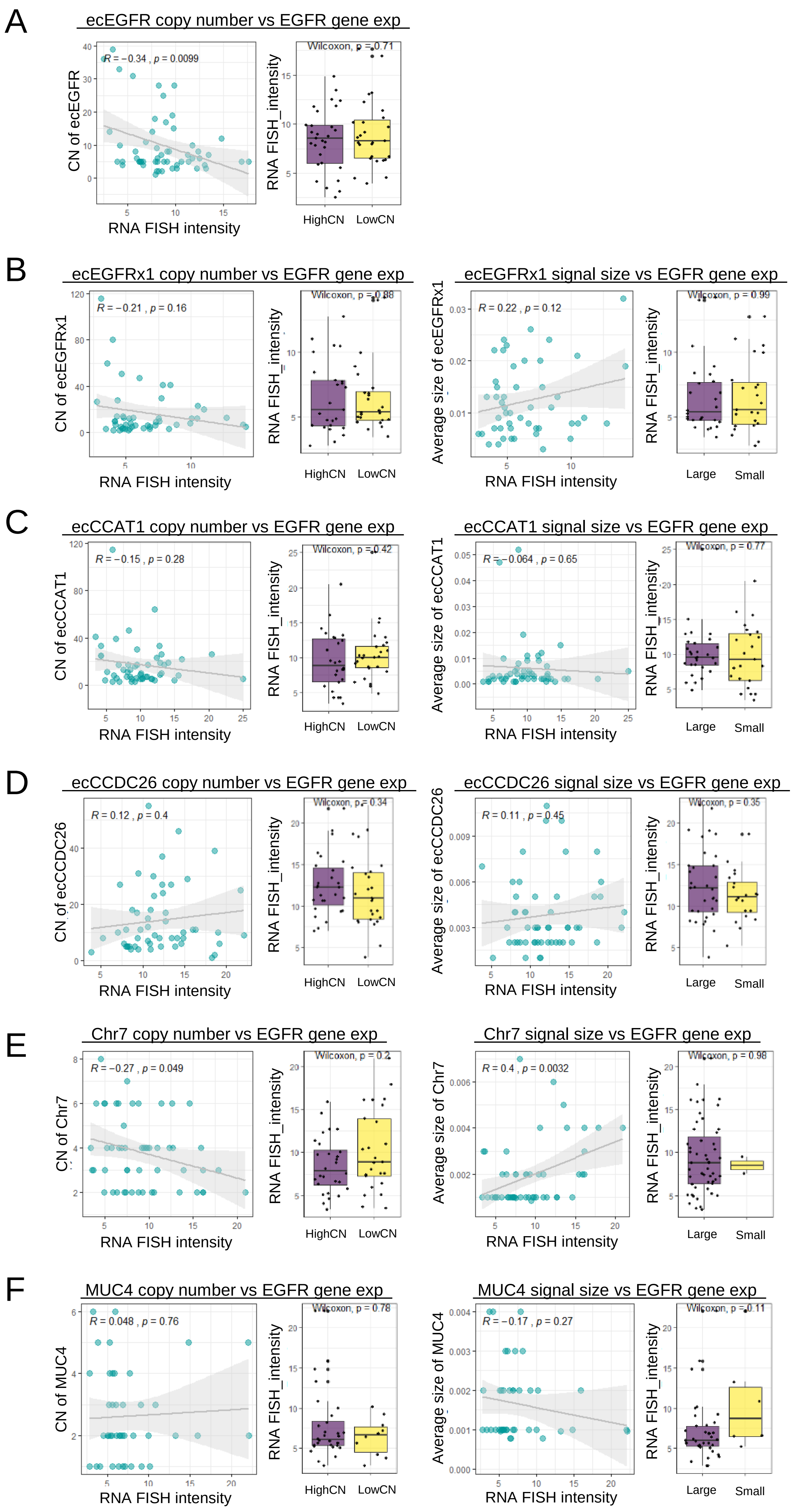

**Supplementary Fig. 9 | Correlation between ecDNA clustering and EGFR gene expression. A.** Correlation between copy number of ecEGFRx1 signals and EGFR gene expression. **B-F.** Correlation between copy number of each Casilio signals and EGFR gene expression (left panels). Correlation between the average signal size of each Casilio signals and EGFR gene expression (right panels). The correlation was determined by Pearson's correlation test. The bar plots represented the comparison of average EGFR gene expression by copy number and signal size (lower panels). The category was determined by its median value. At least 40 single-cell images per group were analyzed.
